# Ca_V_1.2-dependent excitation-transcription coupling modulates nociception

**DOI:** 10.1101/2025.06.25.661480

**Authors:** Jörg Isensee, Patrick Engel, Paul Chu Sin Chung, Chung-Wen Lin, Bushra Saeed Mohammed Dohai, Maximilian Löchte, Maike Siobal, Philipp N. Ostermann, Rebecca Dinnendahl, Mirjam Eberhardt, Inês Carvalheira Arnaut Pombei Stein, Marc R. Suter, Filippo Beleggia, Ruirui Lu, Pascal Falter-Braun, Andreas Leffler, Achim Schmidtko, Tim Hucho

## Abstract

Peripheral nociceptive sensory neurons integrate various noxious inputs, resulting in local depolarization that triggers the firing of action potentials and thus the sensation of pain. We recently reported that nociceptor depolarization itself initiates signaling by the calcium channel Cav1.2 causing acute hyperalgesia *in vivo*. However, whether this mechanism initiates excitation-transcription (E-T) coupling and thereby leads to long-lasting modulation of nociceptor activity remains poorly understood. Using high content imaging of dorsal root ganglion (DRG) neurons, we here found that depolarization of nociceptors induces phosphorylation of the transcription factor (TF) cAMP-response element binding protein (CREB), which was affected by inhibition of protein kinase A (PKA) and calcineurin, but not Ca^2+^/calmodulin-dependent protein kinases. Genetic deletion or pharmacological inhibition of Cav1.2 confirmed its role in calcium-dependent kinase signaling and CREB phosphorylation after depolarization. In line with this, pharmacological modulation of Cav1 channels affected the expression of a subset of depolarization-regulated immediate early genes known to orchestrate a broader transcriptional response. Indeed, RNA-Seq analysis of DRG neurons from mice with a tissue-specific deletion of Cav1.2 in nociceptive sensory neurons (SNS-Cacna1c^-/-^ mice) revealed downregulation of multiple calcium and potassium channel subunits as well as proteins involved in synaptic vesicle release and cell adhesion. Furthermore, repetitive firing of action potentials and release of the neuropeptide CGRP was impaired in Cav1.2-deficient sensory neurons. SNS-Cacna1c^-/-^ mice showed increased sensitivity to noxious heat and exacerbated inflammatory but not neuropathic pain. In conclusion, our data suggest a Cav1.2-dependent E-T coupling mechanism in nociceptors that counteracts nociception *in vivo*.

**Highlights:**

- Depolarization induces Cav1.2- and PKA-dependent phosphorylation of the transcriptional regulator CREB in nociceptors
- Cav1.2 regulates the expression of depolarization-induced immediate early genes and induces reorganization of the signaling network in nociceptors
- Cav1.2-deficient nociceptors downregulate multiple ion channel subunits as well as proteins involved in synaptic vesicle release and cell adhesion
- Cav1.2-deficient nociceptors show impaired depolarization-induced PKA-II signaling, CREB phosphorylation, repetitive firing of action potentials, and release of the neuropeptide CGRP
- Nociceptor-specific Cav1.2-deficiency in mice increases the sensitivity to noxious heat and exacerbates inflammatory pain

**In brief:** Isensee et al. report that depolarization of nociceptors induces calcium influx through Cav1.2 channels, thereby initiating a signaling cascade that links neuronal excitation to transcriptional regulation of immediate early genes (E-T coupling). Nociceptor-specific knockout of Cav1.2 led to transcriptional changes in nociceptor-specific genes and was associated with impaired depolarization-induced signaling, action potential firing, and neuropeptide release. As the Cav1.2-deficient mice show increased noxious heat sensitivity and exacerbated inflammatory pain, these findings suggest that Cav1.2-dependent E-T coupling mechanism counteracts nociception *in vivo*.

## Introduction

Pain is initiated by activation of nociceptive sensory neurons. These sensors of noxious stimuli respond to various physical and chemical inputs that depolarize the neuron locally and eventually trigger the firing of action potentials ^1^. The detection of sensory inputs can undergo dramatic changes in pathologic conditions, a process often referred to as peripheral sensitization ^2,3^. Various mediators then engage receptors to activate intracellular signaling pathways, which include kinases, reactive oxygen and nitrogen species, lipids, and other mediators, followed by the modulation of either noxious sensors (e.g. TRP channels) or ion channels mediating the firing of action potentials (e.g. voltage-gated ion channels) ^4^. In addition to these pathways, nociceptor depolarization itself activates intracellular signaling via the L-type voltage-gated calcium channel Cav1.2, resulting in acute hyperalgesia *in vivo* ^5^. In addition, Cav1-initiated signaling is also a central mechanism for translating acute neuronal activity into transcriptional regulation ^6^. This coupling, often referred to as excitation-transcription (E-T) coupling, has been predominantly investigated in excitatory hippocampal, cortical and sympathetic neurons ^7–10^, but not yet in peripheral nociceptors.

E-T coupling regulates somatic homeostatic processes in neurons, specifically how neurons adapt to changes in activity by decreasing or increasing their overall excitability ^11^. In addition, E-T coupling is essential for synaptic plasticity such as long-term potentiation (LTP) and depression (LTD) ^12,13^. Subsequent to plasticity-inducing stimulation, neurons modulate their synapses using pre-existing macromolecules during the early phase. However, after this period, the formation of lasting LTP/LTD requires de novo gene transcription and protein synthesis in pre- and postsynaptic neurons. Likewise, nociceptor activity potentiates synaptic connectivity to neurons in the spinal dorsal horn ^14,15^. This form of LTP, often referred to as central sensitization, amplifies pain responses in various pain states in animal models and humans ^16,17^. In nociceptors, the mechanisms by which plasticity-inducing stimuli activate intracellular signaling to regulate gene expression remain largely unexplored, and their functional significance is not yet fully understood. The specific mechanism of E-T coupling and its functional outcomes vary in different neuronal subtypes ^6^. In the context of pain, E-T coupling may not only be relevant for central sensitization, but also for the homeostatic regulation of basal nociceptor excitability.

In neurons of the central nervous system, E-T coupling mainly depends on depolarization-induced calcium influx through Cav1, but also Cav2 and *N*-methyl-d-aspartate receptors (NMDARs) are relevant, if calcium from these channels diffuses into Cav1 nanodomains ^9,10,18–22^. Calcium in Cav1 nanodomains binds to calmodulin (CaM) followed by activation of Ca^2+^/calmodulin-dependent protein kinase II (CaMKII) ^9,10,23–25^, which is recruited from docking sites within Cav1 or the β auxiliary subunits ^26–31^. However, in addition to calcium influx, E-T coupling apparently requires a voltage-dependent conformational change of Cav1 and/or its associated proteins ^25^. Cav1 interacts with scaffolding proteins that orchestrate large signalosome complexes ^32^. For instance, the A-kinase interacting protein AKAP79/150 tethers signaling proteins such as protein kinase A (PKA), protein kinase C (PKC), and the CaM-regulated phosphatase calcineurin (CaN) to Cav1 ^33–35^. These associated signaling proteins enable Cav1 channels not only to autoregulate their own activity but also initiate activity-dependent signaling to the nucleus, involving kinases such as CaMKIV ^36^, extracellular signal-regulated kinases (ERK1/2) ^8,37–39^, and PKA ^40–42^.

The activity-dependent signaling response converges on the regulation of TFs such as cAMP response element-binding protein (CREB), serum response factor (SRF), nuclear factor of activated T cells (NFAT), and Myocyte Enhancer Factor 2 (MEF2) ^9,10,36,43–49^. E-T coupling thereby regulates hundreds of activity-induced genes in at least two distinct waves ^50,51^. The first wave results in the transient expression of so-called *immediate early genes* (IEGs), many of which encode TFs ^50^. These IEGs then regulate *late response genes* that define the functional outcome such as the modulation of neuronal excitability^50^. If this is true also in primary afferent nociceptors, is currently unknown. We therefore investigated whether Cav1.2-dependent signaling also mediates E-T coupling in nociceptive neurons and if this has functional relevance for acute and chronic nociceptive pain.

## Results

### Depolarization induces Cav1- and PKA-dependent phosphorylation of the transcriptional regulator CREB in DRG neurons

E-T coupling classically involves Cav1-dependent calcium influx followed by activation of kinase signaling that modulates the activity of transcriptional regulators such as CREB ^47,51–53^. To assess whether ET-coupling is also present in nociceptors, we depolarized rat DRG neurons and quantified phosphorylation of CREB within its kinase-inducible domain at S133 (pCREB). For this, overnight cultures of rat DRG neurons were stimulated with potassium chloride (KCl), fixed, and then labelled for the neuronal marker ubiquitin C-terminal hydrolase L1 (UCHL1), pCREB, and DAPI before being analyzed by High Content (HC) imaging (**Figure 1A**). Automated software-based analysis of > 2000 DRG neurons per time point revealed that KCl-depolarization leads to a rapid and persistent increase in nuclear pCREB, which was not detected after incubation with control solvent (Ctrl) (**Figure 1B**). In line with ET coupling in other neuronal cell models, this increase of nuclear pCREB intensity was absent in the presence of the Cav1 blocker verapamil (**Figure 1B**).

**Figure 1.**
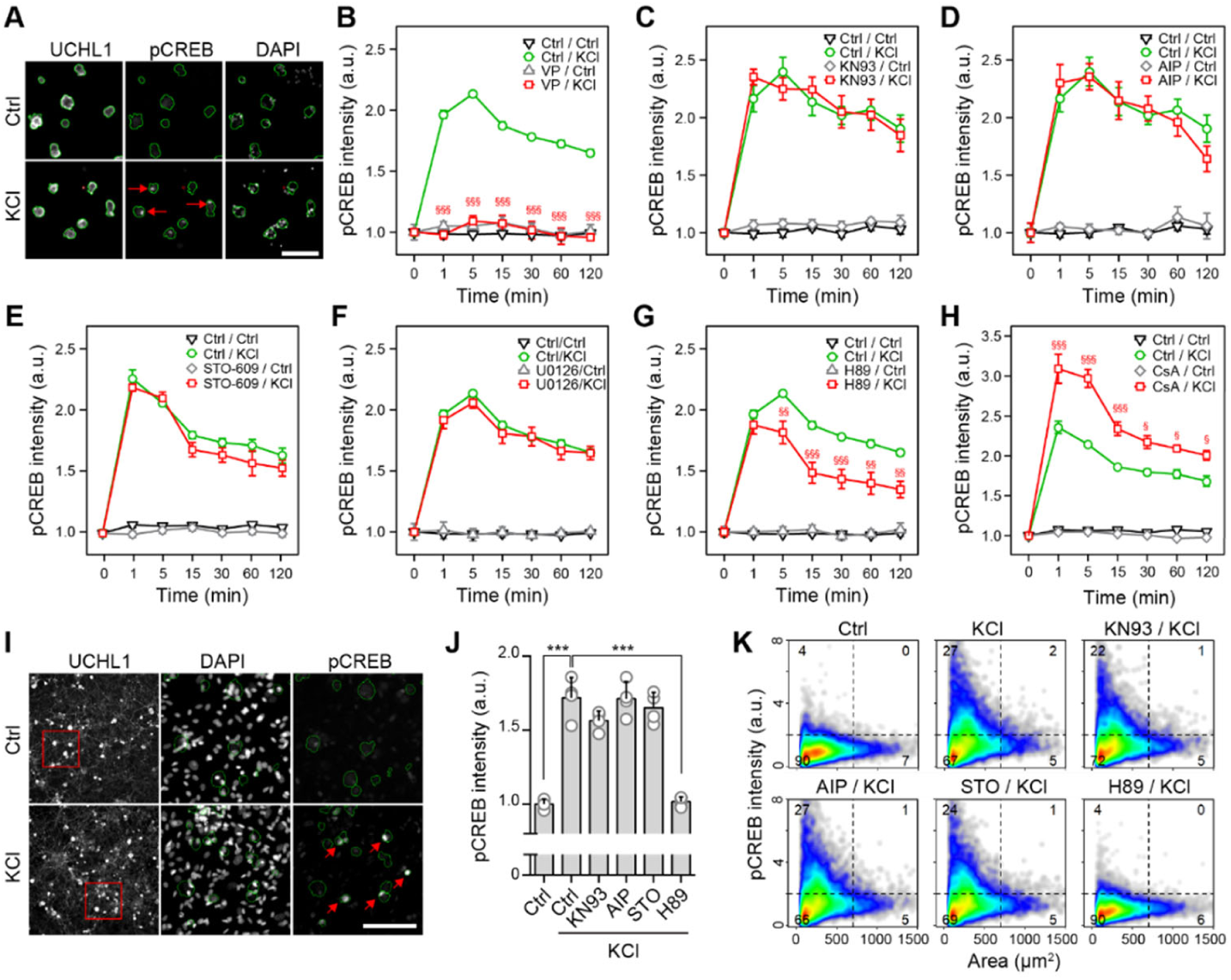
Depolarization induces Cav1- and PKA-dependent phosphorylation of CREB. **(A)** Representative images showing the increase in pCREB immunoreactivity in nuclei of DRG neurons after incubation with 40 mM KCl (red arrows) or control (Ctrl) for 5 min. **(B-H)** Time courses of pCREB immunoreactivity in DRG neurons after incubation with 40 mM KCl or Ctrl, and effects of pre-incubation with the Cav1-inhibitor verapamil (VP, 200 µM, 10 min pre-incubation; **B**), the CaMKII inhibitors KN93 (10 µM, 30 min; **C**) and AIP (1 µM, 30 min; **D**), the CaMKK inhibitor STO-609 (10 µM, 30 min; **E**), the PKA inhibitor H89 (25 µM, 30 min; **F**), and the calcineurin inhibitor cyclosporine A (CsA, 10 µM, 30 min; **G**). **(I)** Representative images of neonatal long-term cultures showing the increase in pCREB immunoreactivity in nuclei of DRG neurons after 5 min depolarization with 40 mM KCl (red arrows) versus Ctrl-stimulated neurons. **(J)** Effect of inhibitors (see above) on the increase in pCREB intensity induced by 5 min stimulation of neonatal DRG cultures with 40 mM KCl. **(K)** Cell density plots showing combined single-cell data of pCREB intensities compared with the cell sizes of the same neurons shown in (J). Dashed lines indicate the gating thresholds used to calculate the percentage of cells in the respective quadrants. Data represent means ± SEM; N = 3-4 experiments with n > 2000 neurons/condition; two-way ANOVA with Bonferroni’s test. \**p* < 0.05; \*\**p* < 0.01; \*\*\**p* < 0.001 indicate significance levels between baseline and stimulated conditions; ^§^*p* < 0.05; ^§§^*p* < 0.01; ^§§§^*p* < 0.001 indicate significance levels between KCl-induced pCREB signals in the absence or presence of (ant-)agonists.

In other neuronal cell models, ET-coupling depends on Cav1-mediated Ca^2+^ influx, Ca^2+^/CaM-dependent activation of α/βCaMKII and shuttling of Ca^2+^/CaM to the nucleus by γCaMKII ^9,24,36,39^. In the nucleus, Ca^2+^/CaM activates CaMKK and its substrate CaMKIV, which finally phosphorylates CREB ^24^. In DRG neurons, however, we did not observe any effect on pCREB by inhibitors of CaMKII (KN93, AIP) or CaMKIV (STO-609) at concentrations two times higher than applied in other studies suggesting that CaMKII and CaMKIV are not part of the Cav1-mediated signaling cascade ^24^ (**Figures 1C-E**).

Calcium influx into neurons is also known to activate the Ras-mitogen-associated protein kinase (MAPK) pathway leading to activation of ERK1/2 ^37^. Activated ERK1/2 phosphorylates and activates ribosomal protein S6 kinase (Rsk), which has been shown to phosphorylate CREB in the nucleus ^21,46,50^. Notably, in a previous study, we observed rapid ERK1/2 phosphorylation (pERK1/2) after KCl-depolarization in DRG neurons that was sensitive to the calcium chelator EGTA as well as Cav1 modulators verapamil and (S)-(-)-Bay K 8644 (BayK), respectively ^5^. However, the MEK inhibitor U0126 did not interfere with pCREB induction in DRG neurons (**Figure 1F**), although U0126 effectively diminished KCl-induced pERK1/2 (**Figure S1**).

Another important protein kinase implicated in depolarization-induced neuronal signaling is protein kinase A (PKA). Early studies suggested that nuclear translocation of ERK1/2 and Rsk2 is mediated by PKA in neurons ^21,46^. In this regard, we have recently reported that calcium in nanodomains near Cav1.2 channels activates the PKA-II isoform in DRG neurons, which modulates Cav1.2 activity by S1928 phosphorylation in a feedback loop^5^. Here, we observed that the PKA inhibitor H89 partially inhibited the pCREB response upon depolarization, which supports a role of PKA-II for E-T coupling in DRG neurons (**Figure 1G**). Dephosphorylation of S1928 by CaN inactivates Cav1 channels ^54,55^. In line with this, pretreatment of DRG neurons with the CaN inhibitor cyclosporine A (CsA) to prevent this dephosphorylation step, reinforced pCREB after depolarization (**Figure 1H**).

While finding central hallmarks of ET coupling such as Cav1- and PKA-dependent CREB phosphorylation, the interconnecting signaling appears to be different in DRG neurons. Much research on ET coupling has been performed on neonatal tissue. To exclude that developmental age and culture conditions underlie the observed differences, we also cultured neonatal DRG neurons for 4 days in conditions similar to those described for neonatal sympathetic neurons ^24^. However, blocking ERK1/2, CaMKII, or CaMKIV did not affect pCREB levels in these neonatal long-term cultures (**Figures 1I-K**), whereas the PKA inhibitor H89 effectively inhibited the pCREB response (**Figures 1J-K**).

### Modulation of Cav1 during depolarization affects the expression of E-T coupling-associated immediate early response genes

Next, we sought to analyze whether excitation of DRG neurons results in alterations of gene expression, thus supporting the existence of E-T coupling in primary afferent nociceptors. We incubated DRG neurons for 6 h with two different concentrations of KCl to induce either weaker (10 mM KCl) or robust depolarization (40 mM KCl) and analyzed changes in gene expression by bulk-RNA-Seq (**Figure 2A, Table S1-2**) ^10,51,56^. In order to facilitate activity of Cav1-channels, we included combinations of the 10 mM KCl concentration with the Cav1-agonist BayK (2 µM). To address the impact of Cav1-channel inhibition, we combined the 40 mM KCl concentration with the Cav1-blocker verapamil (50 µM) ^5^.

**Figure 2.**
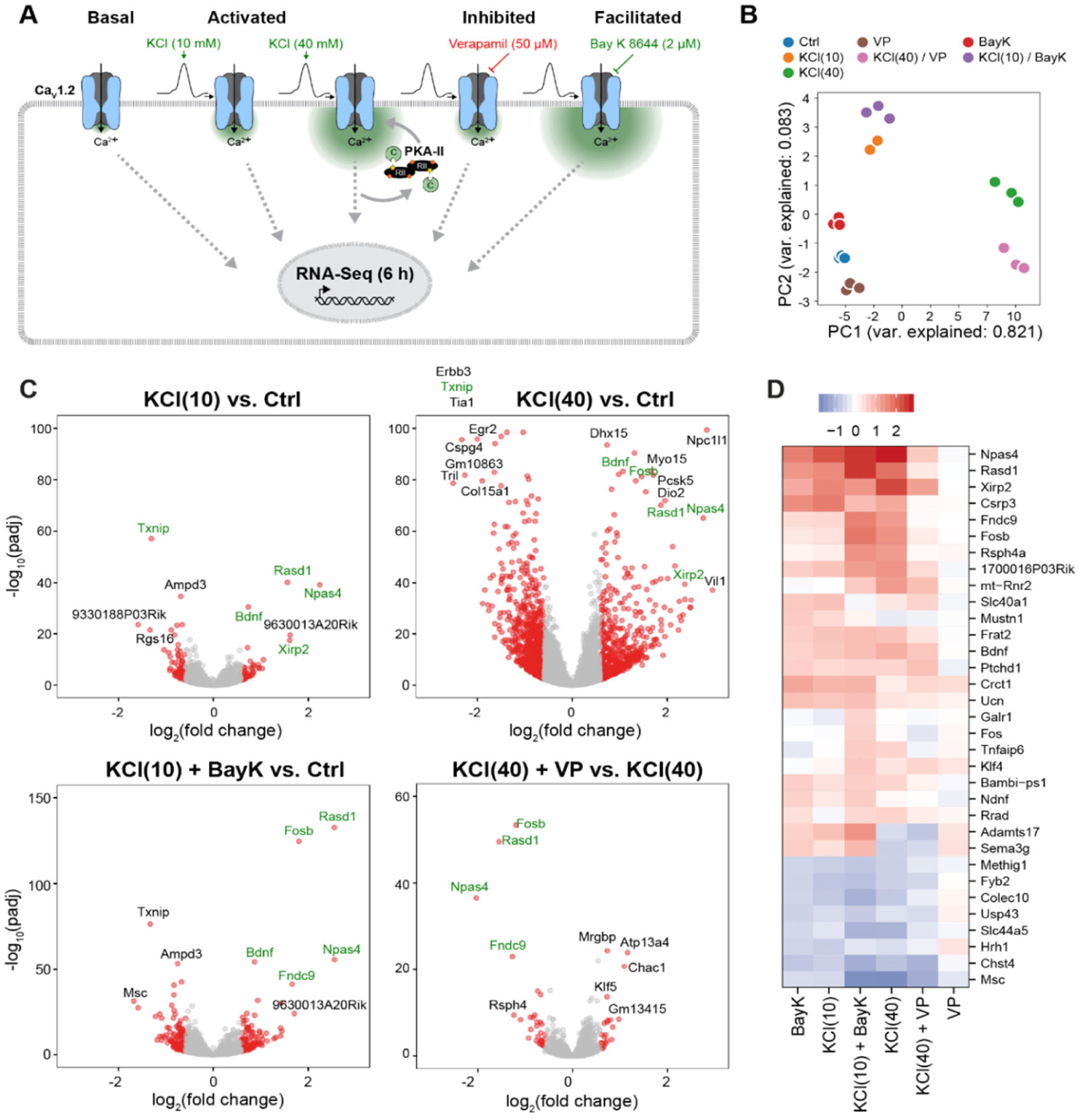
Depolarization leads to Cav1-dependent regulation of gene expression in sensory neurons. **(A)** Experimental design to analyze Cav1-dependent regulation of gene expression. **(B)** Principal component analysis (PCA) of the RNA-Seq data indicating that samples in each group cluster together. **(C)** Volcano plots showing the differentially expressed genes in the indicated comparisons. Red colored dots represent genes with significant adjusted p-values and a fold change >1.5 or <0.667. Several activity-regulated genes are highlighted in green. **(D)** Heatmap of log_2_(fold-changes) of genes showing differential expression (p_adj_ < 0.05 and FC > 1.5-fold) in the comparisons BayK vs. Ctrl and KCl(10)+BayK8644 vs. KCl(10). Data are from n = 3 independent experiments (except condition KCl (10 mM), n=2) with pooled DRG neurons isolated from 14 male C57BL/6N mice per experiment.

KCl-depolarization of the DRG neurons induced concentration-dependent changes in gene expression (**Figures 2B-C**). Of the 21,822 genes detected in our DRG neuron cultures by bulk RNA-seq, stimulation with 10 mM KCl significantly regulated 128 genes (q < 0.05, >1.5-fold change), whereas stimulation with 40 mM KCl affected the expression of 1,563 genes. (**Figures S2A-B**). Overall, the number of differentially expressed genes showing fold-changes > 2-fold is rather small, which may reflect the complex composition of different neuronal and non-neuronal cell types cell in the DRG cultures (**Figure S2A)**. Co-stimulation with BayK with 10 mM KCl nearly doubled the number of differentially expressed genes compared to 10 mM KCl alone (128 vs. 238 genes), supporting the notion that activation of CaV1 channels during depolarization modulates gene expression (**Figures S2A-B**). Applying verapamil to inhibit Cav1 channels did not significantly reduce the number of genes regulated by 40 mM KCl (**Figures S2A-B**). This observation may be explained by: (I) incomplete inhibition of Cav1 channels over the 6-hour period, (II) strong depolarization with 40 mM KCl triggered Cav1-independent transcriptional responses, or (III) verapamil elicited off-target effects that influenced gene expression ^57^. Consistent with Cav1-dependent regulation, several genes such as *Npas4*, *Rasd1*, *Fndc9*, and *Fosb* were upregulated in the presence of the Cav1 agonist BayK but were repressed upon treatment with the Cav1 inhibitor verapamil (**Figure 2C**, highlighted in green). Several of these genes were previously identified as *bona fide* immediate early genes (IEGs) in CNS neurons including *Bdnf*, *Fosb*, *Npas4*, *Rasd1*, *Fundc9*, and *Xirp2* ^51,56^. These IEGs also clustered together in an analysis of all genes either regulated by BayK alone or reinforced by BayK in the presence of 10 mM KCl (**Figure 2D**). Notably, the IEG-encoded TF Npas4 promotes expression of additional IEGs including *Fos*, *Fosb*, *Bdnf*, *Rasd1*, *Crem*, *Cebp*, *Nr4a3*, all of which were also differentially expressed here in DRG neurons after depolarization (**Figures 2C-D**) ^58^. By combining single molecule Fluorescence In Situ Hybridization (smFISH) with immunocytochemical labeling of sensory neurons, we verified the depolarization-dependent regulation of four of five tested genes (*Bdnf*, *Fos*, *Npas4*, *Inhba*, *Rasd1*) in an independent set of samples (**Figure S2C-D**).

### Network analysis suggests E-T coupling changes nociceptive information processing

To gain better insights about functions affected by Cav1-dependent transcription, we analyzed the transcriptomic changes in the context of the protein interaction and signaling network. We constructed a mouse interactome map by combining high-quality mouse and human protein interactions by interolog mapping ^59^. To ensure representation of known pathways, we added mouse KEGG and Reactome pathways to obtain an inferred mouse protein network containing 15,000 nodes and 160,000 edges. Within this network, we defined different types of network neighborhoods for subsequent analyses. In addition to annotated pathways, topological network communities were identified using edge clustering ^60,61^, and, to systematically cover the complete interactome, we included all proteins and their first neighbors in the physical interactome as ‘star communities’. For all interactions we calculated a ‘conditional interaction score’ that captures if the two interactors are regulated in a similar manner. We then aimed to identify network neighborhoods that are coherently regulated, i.e. communities in which the interactions are changed in a similar manner due to Cav1 activity, i.e. a stronger induction following pretreatment with BayK and repression by verapamil. As the transcriptome changes were more specific in the first set of conditions, we used these for module identification, filtering for significantly correlated communities (FDR < 0.05), that show significant differences in regulation (FDR < 0.05, Wilcoxon rank sum test). All of the modules and interactions discussed in the following, are decreased following verapamil treatment.

Top-ranking network modules emerging from this analysis were centered around some, but not all, of the strongly upregulated genes, especially Fosb and Rasd (**Figure S3A**). While some strongly regulated genes (Xirp2 and Npas4) were simply not represented in our network, the neighborhood of Bdnf and Inhba was not notably altered demonstrating the ability of our analysis to identify specific regulated network neighborhoods. Indeed, closer inspection of the found neighborhoods suggests a high relevance for nociception and cellular reprogramming. In several Fosb neighborhoods interactions with Fos, Junb and Jund were most strongly enhanced. Intriguingly, beyond the established role of Fos in E-T coupling in CNS neurons ^50^, a splice isoform of FosB (ΔFosB) has been linked to pain persistence due to chronic stress and inflammation ^62,63^. Intriguingly, the interactions of FosB with multiple effectors of inflammatory pain are strongly enhanced including Ptgs2, encoding COX2, proinflammatory cytokines IL1b and IL6, and the alarmins S100A8 and S100A9 ^64^ indicating that part of the transcriptional response triggers pain-enhancing positive feed-back loop.

More direct indications of signaling network rewiring by Cav1 induced E-T coupling comes from concordantly regulated interactions of Rasd1, which has initially been implicated in inhibition of excitation-secretion (E-S) coupling downstream of corticosteroid signaling ^65^. In fact, the physical star-community around Rasd1 is the most strongly induced neighborhood in the entire network (**Figure S3A**). The interactions of Rasd1 with several gamma subunits of trimeric G-proteins are among the most strongly increased interactions in the entire network thus pointing to possible alterations of GPCR signaling. Intriguingly, the most enhanced interaction in the entire network is with Gngt2, a Gγ subunit shown to inhibit autophagy. The globally second most enhanced interaction pair was Rasd1 interacting with Nr2f6. Knockout of Nr2f6 leads to increased nociception in mice ^66^ and Rasd1 has been described as a negative regulator of Nr2f6 activity ^67^. Intriguingly, the Rasd1 and FosB subnetworks are linked by the catalytic subunit of PKA (Prkaca), which phosphorylates Rasd1 and Fos ^68,69^. Rasd1 can directly inhibit adenylate cyclase ^70^ and cAMP induced gene expression ^71^, thus potentially contributing to termination Cav1 induced transcription.

Finally, we conducted the corresponding analysis for the most decreased interaction pairs. Again, we obtain a large, connected subnetwork but also a smaller network and several disconnected interaction pairs (**Figure S3B**). Intriguingly, corresponding to the prominent position of kinases in the upregulated modules, the decreased subnetworks are centered on a catalytic subunit of protein phosphatase 1 (Ppp1ccb). This large subnetwork is dominated by interactions with other signaling proteins such as protein kinases (Akt3, Camk2d, Camk2g, Prkg1-2, Prkce, Rps6ka3, Rps6ka6, Rock1-2) as well as ion channels such as calcium channels (Cacnb2, Cacna2b1), AMPA-type glutamate receptors (Gria1-4), subunits of the N-methyl-D-aspartate receptor (Grin3a) and the cold sensor Trpm8.

Overall, the network analysis of transcriptomic changes following highly specific pharmacological interventions provides strong evidence for a specific modulation of the transcriptome by Cav1-dependent signaling, which likely affects the information processing in nociceptors. However, given that some of the prior data for the network analysis were obtained in other cellular systems than DRGs, and that the observed changes represent only short-term effects of E-T coupling, it remains difficult to assess the overall impact of these changes on the activity profiles of nociceptors. We therefore turned to an in vivo model.

### Expression of Cav1.2 is reduced in sensory neurons of conditional *Cacna1c* knockout mice

To complementing our pharmacological characterization of Cav1.2-dependent E-T coupling, we generated and validated conditional Cav1.2-deficient mice. DRG neurons mainly express the VGCC α subunits Cav1.2 (L-type), Cav2.1 (P/Q-type), Cav2.2 (N-type), and Cav3.2 (T-type) ^5,72^. Single-cell RNAseq data generated by Zeisel et al. supports expression of the Cav1.2 encoding *Cacna1c* gene in adult peptidergic (PEP1-8) and non-peptidergic nociceptors (NP2-6) but not in mechano- and proprioceptors (NF1-3) or tactile C-low-threshold mechanoreceptors (NP1) of mice (**Figure S4A**)^5,73^. We verified this predominant expression of *Cacna1c* in peptidergic and non-peptidergic nociceptors by re-analyzing the single-cell RNAseq data derived from the independent study by Sharma et al. ^74^ (**Figure S4B**). Importantly, a cross-species transcriptomic atlas of mouse, guinea pig, cynomolgus monkey, and human DRG neurons showed that the expression pattern of *Cacna1c* is conserved between mice and humans ^75^ (**Figure S4C**).

Expression of the Nav1.8 encoding *Scn10a* gene shows considerable overlap with *Cacna1c* (**Figure 3A**). To generate conditional *Cacna1c*-deficient mice, we crossed *Cacna1c^fl^*^/fl^ mice ^76^ with mice expressing Cre recombinase under control of the sensory neuron-specific (SNS) *Scn10a* gene promoter ^77^. Notably, Cre-mediated recombination begins perinatally in *Scn10*^Cre/+^ driver mice and therefore should not affect embryonic development ^77^. To validate the conditional *Cacna1c* knockout on RNA level, we analyzed bulk RNA derived from DRGs. This analysis verified a substantial reduction of *Cacna1c* mRNA by 75.8% in SNS-Cacna1c^-/-^ mice compared to floxed control littermates (12.9 ± 1.6 vs. 3.1 ± 0.2 TPM, n = 3 mice, **Figure 3B**). To analyze the *Cacna1c* knockout on protein level, we performed HC imaging of Cav1.2 immuno-labelled DRG neurons (**Figure 3C**). Since the Cav1.2-specific antibody produced weak signals even if using long exposure times, we subtracted the mean background signal determined by omitting the primary antibody (**Figure 2C**, blue line). We observed that the mean Cav1.2 intensity was reduced by 25.8 % in the SNS-Cacna1c^-/-^ DRG neurons, which verifies the Cav1.2 knockout at protein level (**Figure 3D**).

**Figure 3.**
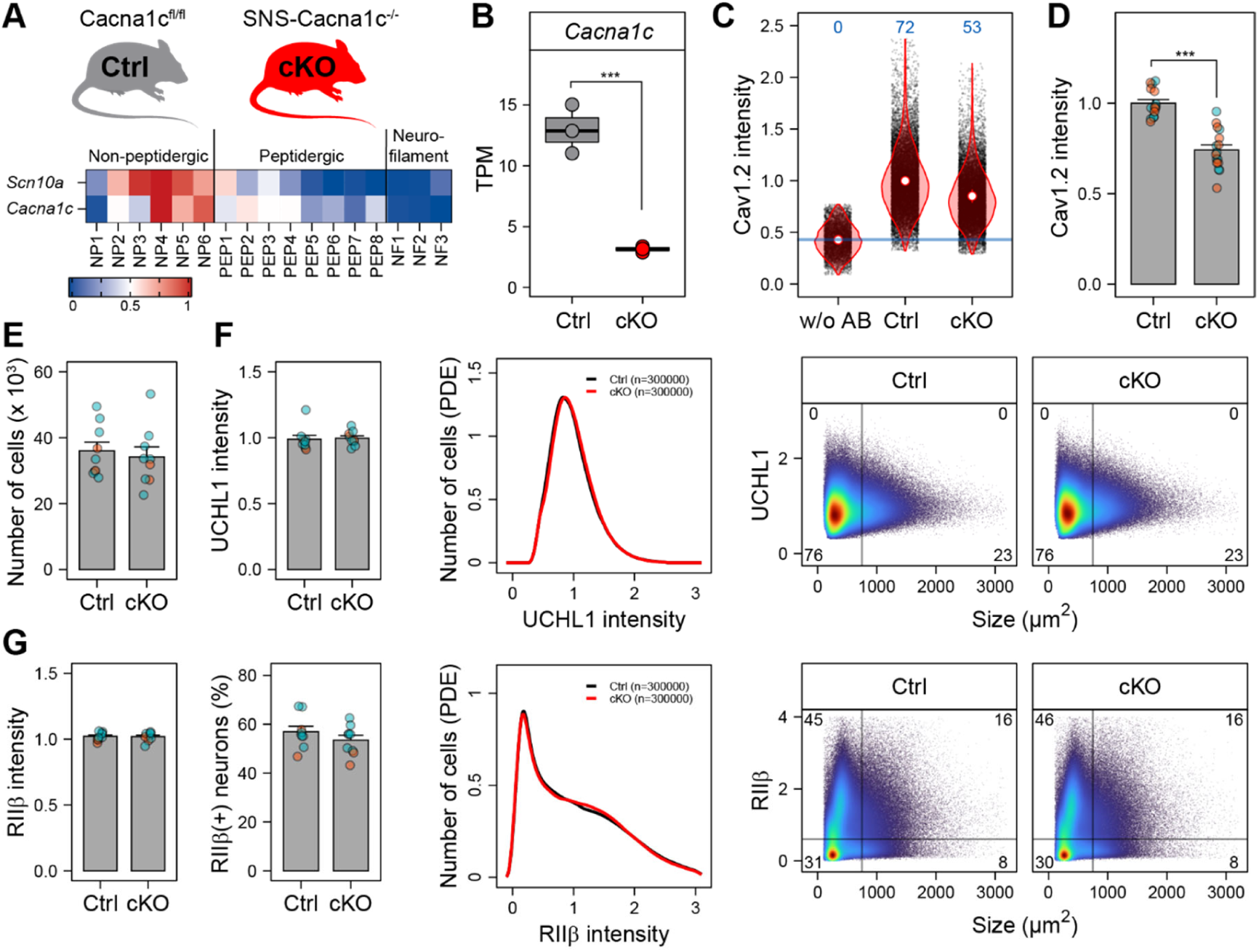
Expression of Cav1.2 is reduced in sensory neurons of conditional Cacna1c knockout mice. **(A)** Conditional Cav1.2 knockout mice (cKO, SNS-Cacna1c^-/-^) were generated by crossing Cacna1c^fl/fl^ (Ctrl) mice with mice expressing Cre under control of the sensory neuron-specific (SNS) *Scn10a* gene promoter. Single cell RNA-Seq data ^73^ indicate that *Scn10a* encoding Nav1.8 is coexpressed with *Cacna1c* in peptidergic (PEP) and non-peptidergic nociceptors (NP), but not neurofilament-expressing mechano- and proprioceptors (NF). **(B)** Bulk RNA-Seq results for *Cacna1c* mRNA of DRGs from Cacna1c^fl/fl^ versus SNS-Cacna1c^-/-^ mice. TPM; transcripts per million. **(C)** HC imaging single cell data of DRG neurons indicating downregulation of Cav1.2 protein in neurons of SNS-Cacna1c^-/-^ mice. **(D)** Mean values of replicate wells form n = 4 (2 male, 2 female) mice of the data shown in (C). Values of males are colored in blue, females in red. **(E)** Number of analyzable DRG neurons per isolation from n = 9 female (red) and male (blue) mice per genotype. **(F-G)** UCHL1 and RIIβ expression pattern in N = 9 mice (female in red and male in blue) of n = 300.000 DRG neurons of per genotype.

### Cav1.2-deficiency does not affect the composition of sensory neuron subgroups

During mammalian development, electrical activity promotes the calcium-dependent survival of neurons with appropriate synaptic connections ^47^. Expression of both, Nav1.8 and Cav1.2, starts before E15 in the late embryonic development of rodents, and reaches adult levels in early postnatal periods ^74,78–80^. Since Cav1.2 is important for E-T coupling, neuronal survival, and synaptic efficacy, we worried that Cav1.2-deficiency leads to changes in the number and composition of neuronal subgroups ^81^. To characterize the SNS-Cacna1c^-/-^mice in this regard before conducting functional assays, we determined the number of viable neurons and analyzed the expression patterns of the sensory neuron marker UCHL1 and the nociceptor marker RIIβ in >300.000 DRG neurons derived from mice of both genotypes (**Figures 3E-G**). Furthermore, we analyzed marker protein expression for non-peptidergic subgroups (CaMKIIα, IB4, Nav1.8), peptidergic subgroups (TRPV1), and myelinated proprio- and mechanoreceptors (NF200) in >14.000 neurons per genotype (**Figure S5**). Overall, this analysis did not reveal any alterations in the composition and abundance of sensory neuron subgroups in SNS-Cacna1c^-/-^ DRGs.

### Cav1.2-deficiency reduces calcium influx into DRG neurons

Combining genetic and pharmacological targeting allows to define the role of Cav1.2 in more detail. First, we investigated how much of the depolarization-induced calcium influx depends on Cav1.2. In overnight cultures of DRG neurons loaded with the far-red calcium imaging dye Calbryte 630 AM (**Figures 4A-B**), strong KCl-depolarization (40 mM) evoked a similar response in SNS-Cacna1c^-/-^ neurons compared to control neurons (**Figures 4C-D**, Ctrl). Thus, Cav1.2 deficiency does not affect the overall calcium influx into DRG neurons upon strong depolarization. However, mild depolarization with 10 mM KCl resulted in significantly less calcium influx in Cav1.2-deficient neurons (**Figures 4C-D**, KCl(10)). Thus, Cav1.2-deficient neurons exhibit an altered rate of calcium influx in specific conditions. At early time points, pre-treatment with the Cav1 agonist BayK increased the rate of calcium influx mainly in control neurons, which supports the Cav1.2 conditional knockout in SNS-Cacna1c^-/-^ mice (**Figure 4C**, KCl(10) vs. BayK + KCl(10)). In contrast, calcium influx induced by the VGSC agonist veratridine (VT) or the TRPV1 agonist capsaicin (Cap) was not altered in Cav1.2-deficient neurons (**Figures 4C-D**, VT and Cap). Thus, our data corroborated that Cav1 channels open at mild depolarization in neurons ^82^, although they were initially classified as ‘high-voltage activated’.

**Figure 4.**
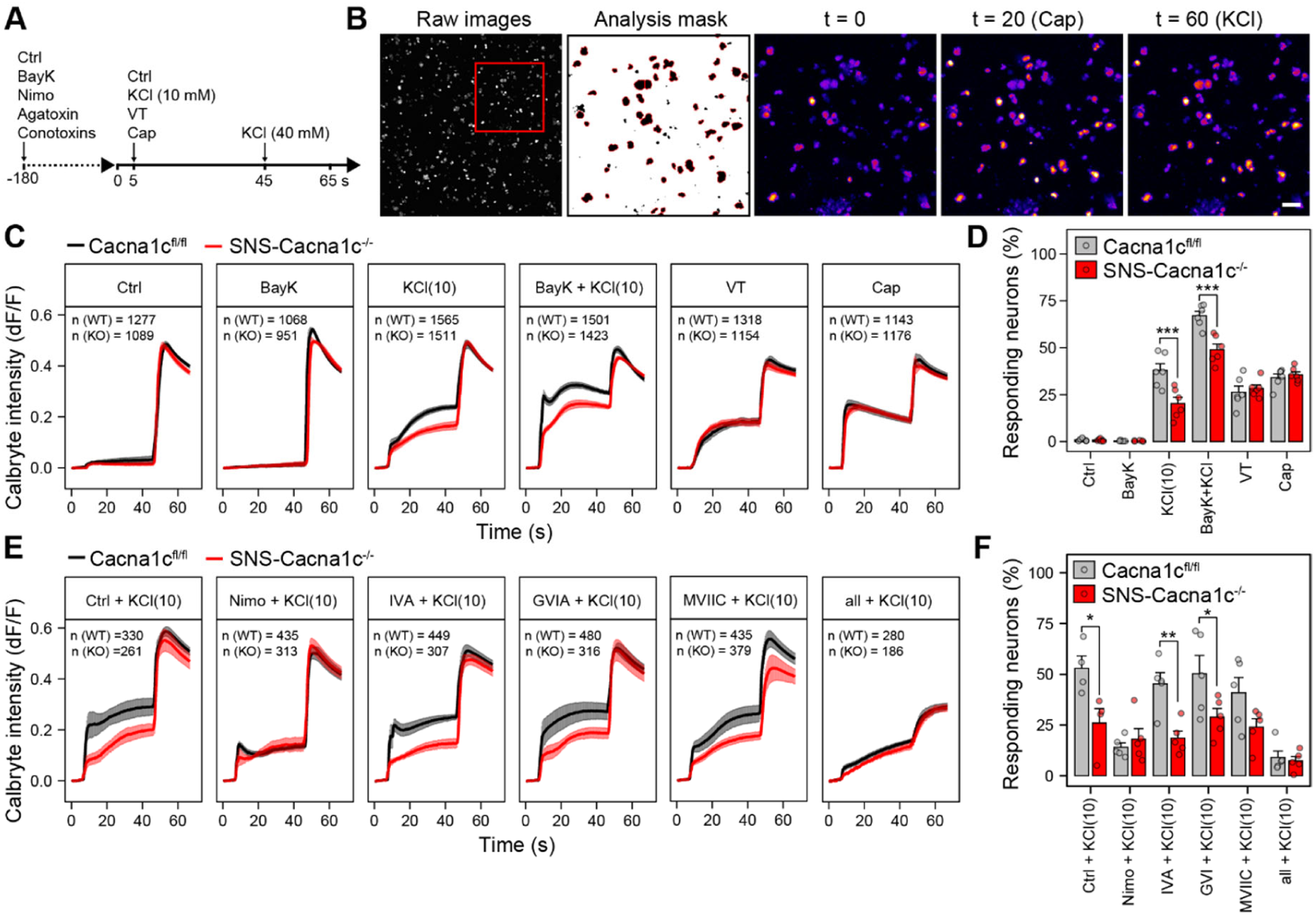
Cav1.2-deficiency decreases calcium influx after depolarization of DRG neurons. **(A)** Overview of the time course to analyze calcium influx using Calbryte 630 dye. **(B)** Representative example images and analysis mask used to quantify the Calbryte 630 fluorescence intensity in DRG neurons. Scale bar, 100 µm **(C)** Mean traces after depolarization with a low dose of KCl (10 mM) in the absence or presence of the Cav1 agonist (S)-(-)-Bay K 8644 (BayK) as well as with veratridine (VT, 100 µM) or capsaicin (Cap, 10 µM). N = 6 experiments, n = 951-1565 neurons **(D)** Mean percentage (15-40 s after stimulation) of responding neurons in experiments shown in C. **(E)** Mean traces after depolarization with a low dose of KCl (10 mM) in the absence or presence of the Cav inhibitors nimodipine (Nimo, 10 µM), ω-agatoxin IVA (ω-AGA IVA, 1 µM), ω-conotoxin GVIA (ω-CTX GVIA, 3 µM), ω-conotoxin MVIIC (ω-CTX MVIIC, 1 µM), or a combination of all inhibitors (All inh.). N = 4-5 experiments with n = 186-480 neurons. **(F)** Mean percentage (15-40 s after stimulation) of responding neurons in the experiments shown in E. Data represent means ± SEM; two-way ANOVA with Bonferroni’s test. \**p* < 0.05; \*\**p* < 0.01; \*\*\**p* < 0.001.

To further corroborate the role of Cav1.2 for calcium influx after mild depolarization, we then established pharmacological conditions for the specific inhibition of Cav1.2 in DRG neurons. Structurally unrelated pore blockers such as phenylalkylamines (i.e. verapamil) and benzothiazepines (i.e. diltiazem) are effective, but suffer from non-specific effects ^83,84^. In contrast, 1,4-dihydropyridine-based (DHP) allosteric modulators such as nimodipine are highly specific, but are state-dependent and often do not completely inhibit Cav1 channel function in neurons ^82,85^. We therefore investigated the effect of nimodipine and verapamil on calcium influx after mild (10 mM) and strong (40 mM) KCl-depolarization (**Figures 4E-F**). Both inhibitors more efficiently suppressed calcium influx after mild depolarization of control neurons compared to Cav1.2-deficient neurons supporting that mild depolarization opens Cav1.2 (**Figures 4E-F and S6**).

Since Cav1.2-deficiency only affects calcium influx after mild depolarization, we were wondering whether Cav2 channels (Cav2.1, Cav2.2) mediate the calcium influx during strong depolarization. Therefore, we administered high doses of toxins to inhibit Cav2.1 (ω-agatoxin IVA) and Cav2.2 (ω-conotoxin GVIA) separately or simultaneously (ω-conotoxin MVIIC). These toxins did not show substantial inhibitory effects in both genotypes if applied alone (**Figures 4E-F**). However, combining the toxins with nimodipine led to substantially reduced calcium influx after mild and strong depolarization (**Figures 4E-F**).

### Cav1.2-deficiency decreases depolarization-induced PKA-II and CREB activity

We have shown that Cav1.2-mediated signaling activates PKA-II ^5^, which contributes to phosphorylation of CREB driving E-T coupling in DRG neurons (**Figures 1,2**). To confirm this data in Cav1.2-deficient mice, we depolarized the DRG neurons with increasing concentrations of KCl (0-80 mM) and analyzed the fixed cells by HC imaging. Although PKA has been indirectly implicated with E-T coupling in neurons ^40–42^, its activity could not be studied directly at the cellular level. Here, we applied an antibody that binds to the phosphorylated inhibitory sites of PKA-II regulatory subunits RIIα and RIIβ (collectively referred to as pRII) only in dissociated, active PKA-II kinases ^86^, which allows the quantification of endogenous PKA-II activity in UCHL1-positive DRG neurons ^5,86–88^. To estimate whether the response occurs in nociceptors, we also quantified the expression level of the PKA regulatory subunit RIIβ, a marker expressed in most nociceptive DRG neuron subgroups ^87^. We observed a significant right shift of the dose response curve as well as a reduction of the amplitude of the KCl-induced PKA-II activity (pRII) in Cav1.2-deficient neurons (Figure 5A). In the presence of the Cav1 agonist BayK, the right shift of the dose response curve was even more pronounced (**Figure 5B, E**). This reduction of PKA-II activity in Cav1.2-deficient neurons was greater in the RIIβ(+) neuron population indicating that nociceptors are more affected (**Figures S7A-D**). Intriguingly, BayK also reduced the amplitude of pRII induction, which was absent in Cav1.2-deficient neurons (**Figures 5C-D**). This inhibitory effect of BayK likely represents an antagonistic activity of the compound. BayK is an agonistic stereoisomer of the DHP antagonist (R)- (+)-BayK8644 and therefore structurally similar to other DHP antagonists such as nimodipine ^89^. BayK is known to act as an antagonist when applied at higher concentrations ^89,90^.

**Figure 5.**
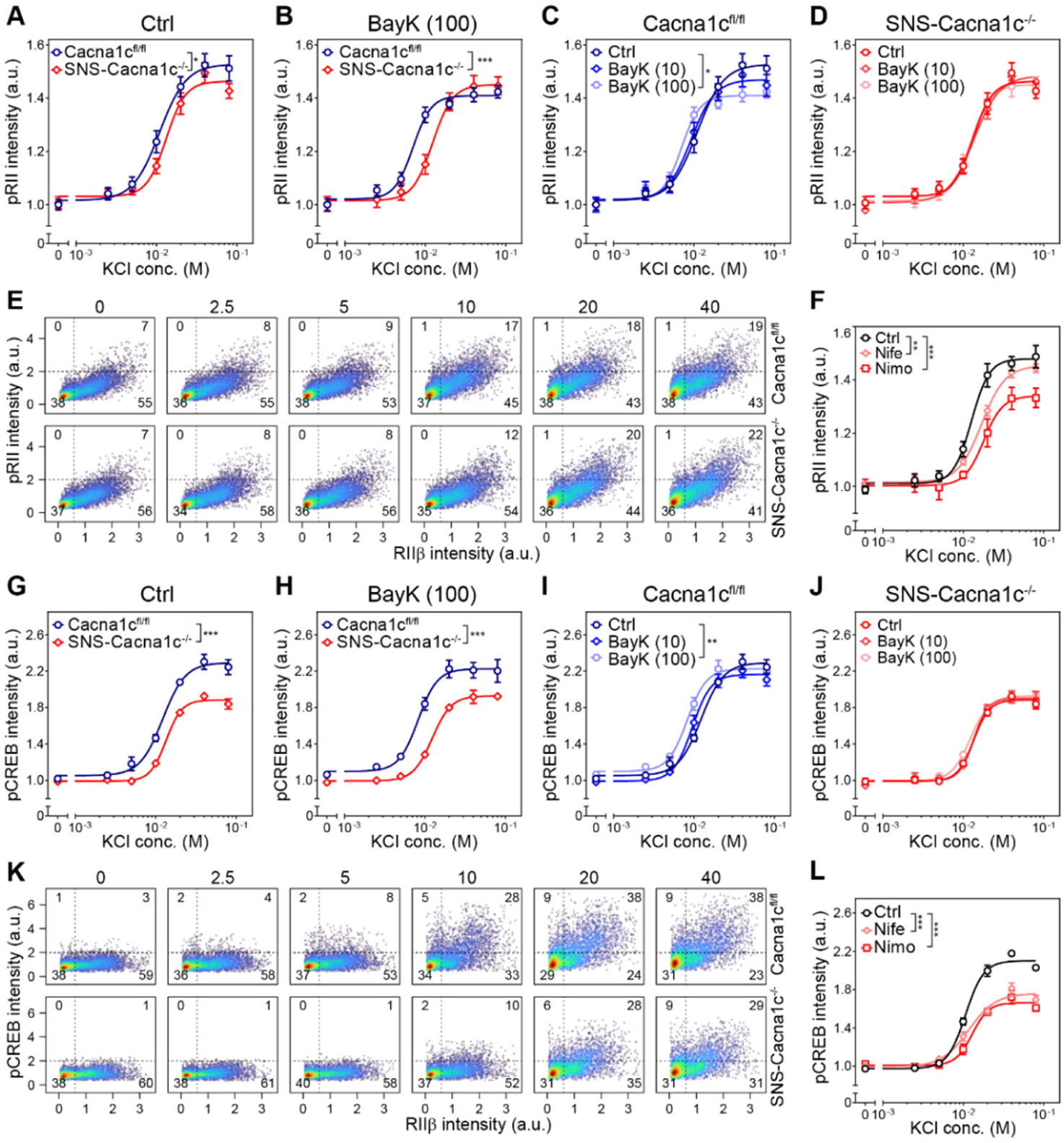
Cav1.2-deficiency decreases depolarization-induced PKA-II and CREB activity. **(A-D)** Concentration-response curves of pRII intensities in DRG neurons of Cacna1c^fl/fl^ and SNS-Cacna1c^-/-^ mice depolarized with KCl (0-80 mM) for 3 min. Neurons in B-D were preincubated with BayK (10 or 100 nM) for 10 min. **(E)** Cell density plots of single cell data shown in B. Dashed lines indicate gating thresholds to discriminate between RIIβ(−) and RIIβ(+) neurons with the numbers indicating the relative percentage of cells in the respective quadrant. **(F)** Concentration-response curves showing the effect nimodipine or nifedipine (10 µM, 10 min) on pRII signals induced by KCl-depolarization. **(G-J)** Dose response curves of pCREB intensities in DRG neurons of Cacna1c^fl/fl^ and SNS-Cacna1c^-/-^ mice depolarized with KCl (0-80 mM) for 3 min. Neurons in H-J were preincubated with BayK (10 or 100 nM) for 10 min. **(E)** Cell density plots of combined single cell data shown in H. Dashed lines indicate the gating thresholds used to calculate the percentage of cells in the respective quadrants. **(L)** Dose-response curves showing the effect nimodipine or nifedipine (10 µM, 10 min) on pRII signals induced by KCl-depolarization. Mean values of RIIβ(-) vs. RIIβ(+) neurons are shown in Fig S5. Data are means ± SEM; N = 3-4 experiments with a total of n > 2000 neurons/condition; extra-sum-of-squares F test; \**p* < 0.05; \*\**p* < 0.01; \*\*\**p* < 0.001 indicate significance levels between genotypes or stimulated conditions.

Similar to Cav1.2-deficiency, preincubation with nimodipine or nifedipine led to partial inhibition of the PKA-II response (**Figure 5F**). This effect of nimodipine was absent in Cav1.2-deficient DRG neurons, indicating specificity (**Figures S7M-N**). Since pharmacological inhibition or even knockout of Cav1.2 only slightly reduced PKA-II activation, we hypothesized that calcium influx via other VGCCs might be sufficient to activate PKA-II at stronger depolarization. We therefore tested for the effect of the Cav2 antagonists ω-agatoxin IVA (Cav2.1), ω-conotoxin GVIA (Cav2.2), and ω-conotoxin MVIIC (Cav2.x) alone or in combination with nimodipine. The toxins did not show substantial inhibitory effects on their own (**Figures S8A-C**), even if all toxins were combined (**Figure S8D**). However, co-administering the toxins with nimodipine completely suppressed PKA-II activation, suggesting that calcium influx through both Cav1 and Cav2 channels is sufficient to activate PKA-II (**Figure S8D**).

PKA-II, in concert with other kinases, regulates the activity of CREB in the nucleus, which is a key mechanism of E-T coupling. In Cav1.2-deficient neurons, we observed a rightward shift of the dose response and a 29% lower maximal pCREB amplitude (**Figure 5G**). Again, the presence of the Cav1.2 agonist BayK led to a leftward shift and reduction of the maximal amplitude in the control neurons, but not in the Cav1.2-deficient neurons (**Figures 5H-K**). Resembling the effect of genetic Cav1.2 knockout, preincubation with the nimodipine or nifedipine led to similar inhibition of pCREB after depolarization (**Figure 5L vs 5G**). Moreover, this inhibitory effect of nimodipine was almost absent in DRG neurons of SNS-Cacna1c^-/-^ mice (**Figures S5O-P**). Overall, these findings substantiate that Cav1.2 signaling contributes to PKA-II activation and CREB phosphorylation in DRG neurons.

### Cav1.2-deficiency leads to differential gene expression in DRG neurons *in vivo*

The conditional knockout model now allowed us to further investigate Cav1.2-dependent gene expression *in vivo* and in absence of acute stimulation. By analyzing the transcriptome of whole DRGs from SNS-Cacna1c^-/-^ and control littermates using bulk RNA-Seq, we identified 437 differentially expressed protein-coding genes in the SNS-Cacna1c^-/-^ model (FC > 1.5-fold, padj < 0.05), most of which were downregulated (320 down-vs 117 upregulated genes) (**Figure 6A**). As expected by the knockout only in Nav1.8-expressing neurons, Cav1.2 expression levels were significantly reduced to 23% in the SNS-Cav1.2^-/-^ mice as compared to control littermates (**Figures 2B and 6B-C**, red gene symbol).

**Figure 6.**
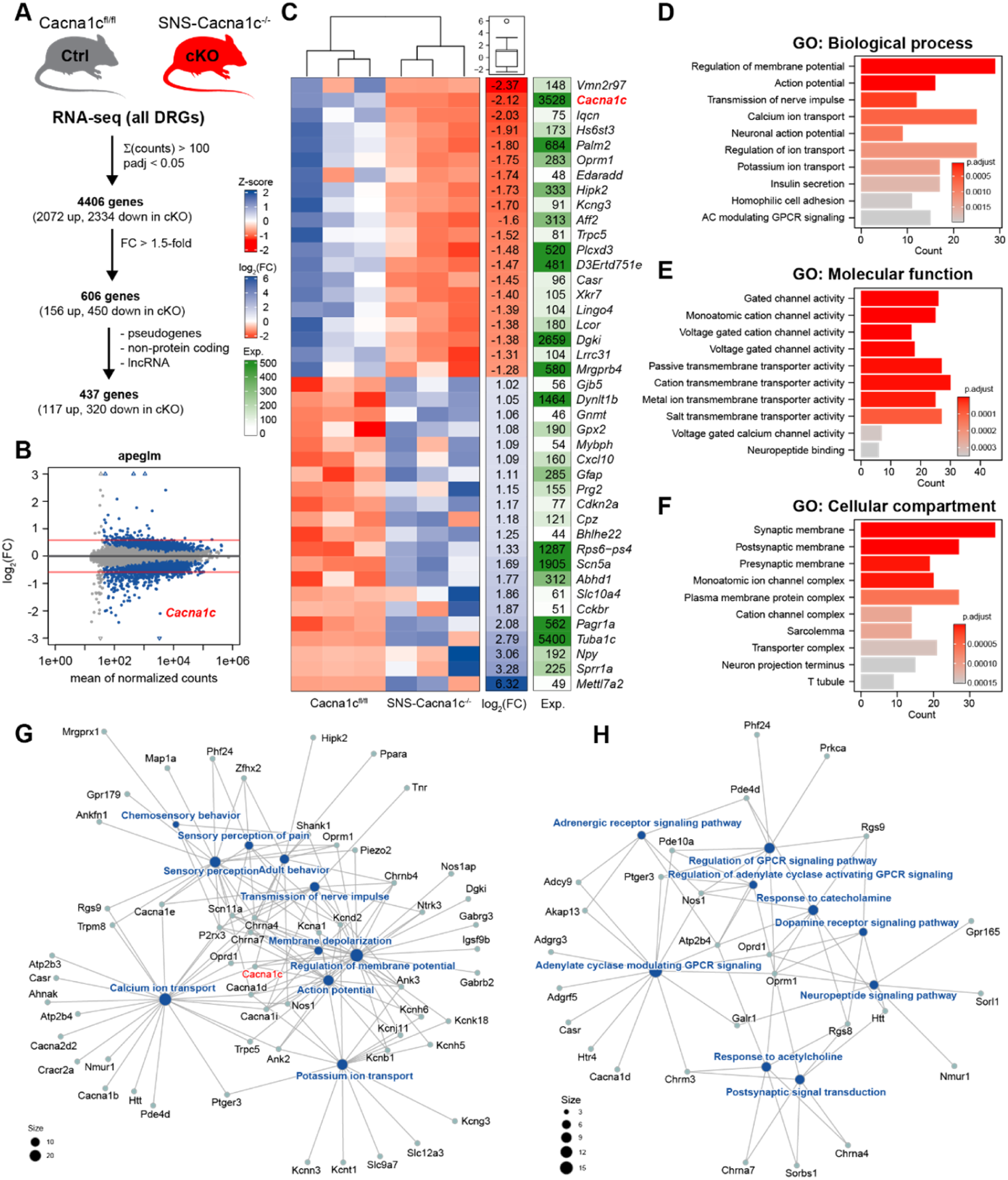
Cav1.2-dependent E-T coupling induces transcriptional regulation *in vivo*. **(A)** Experimental design and numbers of differentially expressed genes in DRGs of SNS-Cacna1c^-/-^ versus Cacna1c^fl/fl^ mice. **(B)** MA plot showing the log_2_(fold-change) versus the normalized counts. For visualization, the log_2_(fold-change) was estimated using the empirical Bayes shrinkage estimation algorithm apglm ^122^. Blue spots indicate genes with significant differential expression (*p*_adj_ < 0.05). **(C)** Heatmap of the normalized and scaled counts of the top 20 differentially expressed genes. The mean of the log_2_(FC) and counts (Exp.) are shown on the right. **(D-F)** Top 10 Gene Ontology (GO) terms for biological process, molecular function, and cellular compartment identified using genes downregulated in DRGs of SNS-Cacna1c^-/-^ versus Cacna1c^fl/fl^ mice (log2(FC) > 1.5-fold, *p*_adj_ < 0.05). **(G-H)** Network plot of enriched terms and associated genes related to action potential firing (G) and GPCR signaling (H).

To analyze the gene ontologies (GO) associated with the identified differentially expressed genes, we divided the dataset in up- and downregulated gene sets. Among the upregulated genes, we did not detect any associated functional terms with p values < 0.001. In contrast, the downregulated genes were significantly enriched for functional terms related to neuronal activity and synaptic functions such as action/membrane potential, calcium and potassium ion transport, GPCR signaling, neuropeptide secretion, vesicle fusion, and adhesion (**Figures 6D-E** and **Table S2**). With regard to the affected cellular compartment, mainly GO terms related to the pre- and postsynaptic membrane and ion channel complexes were found (**Figure 6F**). Hierarchical clustering based on the pairwise similarities of the enriched biological processes revealed six groups of similar terms related to action potential generation and sensory perception, adhesion, learning, GPCR signaling, and secretion (**Figure S9**). Since calcium-influx through Cav1 channels is also important for muscle contraction, many terms related to muscle physiology have also been identified.

Notably, although Cav1.2-deficiency has been shown to result in compensatory upregulation of related Cav1 isoforms ^91^, we observed the downregulation of multiple other calcium channel subunits and associated regulatory proteins in SNS-Cacna1c^-/-^ mice (**Figure 6G**). In particular, the α-subunits *Cacna1b* (Cav2.2) as well the lower expressed *Cacna1d* (Cav1.3), *Cacna1e* (Cav2.3), and *Cacna1i* (Cav3.3) were downregulated. In addition, also the associated α2δ2 subunit (*Cacna2d2*) and the calcium-transporting ATPase 4 (*Atp2b4*) was lower expressed in neurons from SNS-Cacna1c^-/-^ mice. Multiple potassium channel subunits such as *Kcna1* (K_V_1.1), *Kcnb1* (K_V_2.1), *Kcnd2* (K_V_4.2), *Kcng3* (K_V_6.3), *Kcnh5* (K_V_10.2), *Kcnh6* (K_V_11.2), *Kcnj11* (Kir6.2), *Kcnk18* (TRESK), *Kcnn3* (KCa2.3), and *Kcnt1* (Slack) were also downregulated.

According to respective databases (OMIM, Pain Genes DB) and literature, we identified multiple genes with crucial functions for nociception and nociceptor physiology (**Table S3**). For instance, several GPCRs known to regulate nociceptor sensitization including the µ and δ-opioid receptors *Oprm1* and *Oprd1*, the purinergic receptor *P2rx3*, the prostanoid receptor *Ptger3*, the neuropeptide receptors *Galr1* and *Nmur1*, as well as several Mas-related GPCRs such as *Mrgprb4*, *Mrgprd*, and *Mrgprx1* were differentially expressed (**Figure 6H**). Ion channels directly involved in temperature detection (*Trpm8*), mechanosensation (*Piezo2*), and neuronal excitability (*Scn11a*) were differentially expressed as well. Furthermore, the putative transcriptional regulator *Zfhx2* was downregulated in SNS-Cacna1c^-/-^ mice. LOF-mutations in *Zfhx2* induce hyposensitivity to noxious heat and painless bone fractures, which has been confirmed in a knockout mouse model ^92^. Among the rather few neuronal genes being upregulated in SNS-Cacna1c^-/-^ DRGs, we identified the neuropeptide Y (Npy), the CGRP receptor *Ramp1*, as well as the cardiac TTX-sensitive VGSC *Scn5a*. However, in comparison to other DRG-specific VGSCs such as Nav1.7-9 with TPM values of 50-200, expression levels of *Scn5a* were rather low (TPM = 2.8 and 8 in Cacna1c^fl/fl^ and SNS-Cacn1c^-/-^ mice).

Overall, the RNA-Seq analysis confirmed that loss of Cav1.2 leads to altered expression of neuronal genes in DRGs *in vivo* without applying any experimental intervention. The basal neuronal activity of the sensory nervous system is probably sufficient to induce Cav1.2-dependent E-T coupling, suggesting that this mechanism regulates basal electrical activity of DRG neurons. Cav1.2-deficiency leads to complex adaptations that affect calcium, potassium, and sodium homeostasis, as well as associated signaling mechanisms.

### Cav1.2-deficiency reduces repetitive firing of DRG neurons

By RNA-Seq, we identified multiple ion channels with altered expression levels in the DRG neurons of SNS-Cacna1c^-/-^ mice. To test whether this has significant impact on evoked calcium currents (ICa), we performed whole-cell patch-clamp recordings with small-diameter DRG neurons. Currents were elicited by applying 100-ms-long test pulses in steps of of 10 mV from a holding potential of -90 mV. In line with the literature, we observed a large variability of calcium transients among the tested neurons (**Figure 7A**) ^93^. The current density remained largely unaffected in neurons of SNS-Cacna1c^-/-^ mice, although a trend towards lower values was observed (**Figure 7B**). This is in line with the calcium imaging experiments (**Figure 3**) showing that only a smaller proportion of the total calcium influx is mediated by Cav1.2 in DRG neurons.

**Figure 7.**
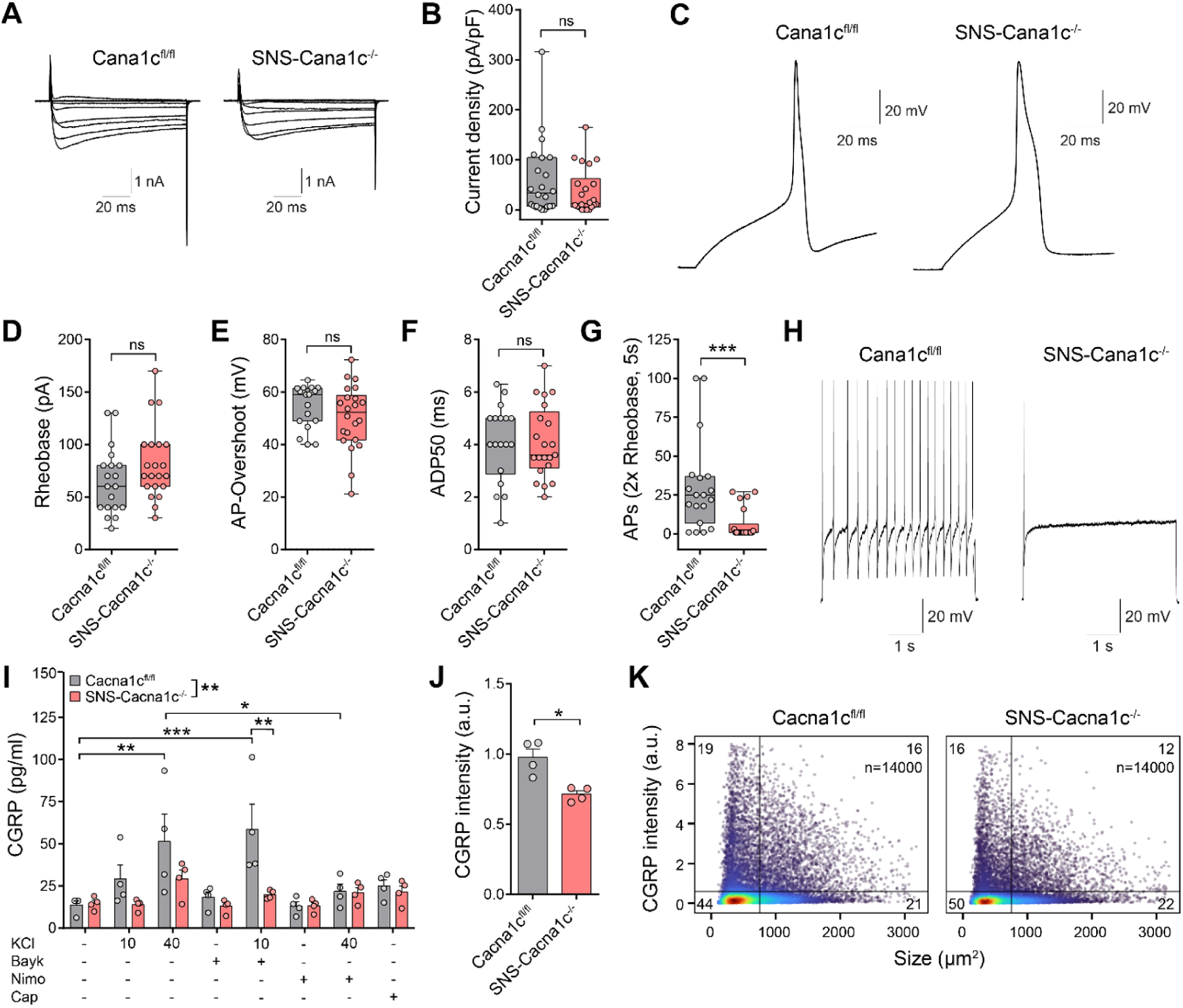
Cav1.2 deletion decreases repetitive firing and CGRP release in DRG neurons. **(A)** Representative traces of calcium currents (ICa) in DRG neurons from Cacna1c^fl/fl^ and SNS-Cacna1c^-/-^ mice measured by whole-cell patch-clamp recording. **(B)** ICa current density (n = 22; N = 3 mice per group). **(C)** Representative AP traces of whole-cell patch-clamp recordings. **(D)** Minimal current to elicit an AP (Rheobase) (n = 19-22; N = 3 mice per group) **(E)** AP-Overshoot (n = 19-22; N = 3 mice per group). **(F)** Duration of 50% repolarization after AP firing (ADP50) (n = 18-21; N = 3 mice per group). **(G)** Number of action potentials (APs) within 5 s evoked by intracellular current injection (2x Rheobase) (n = 19-22; N = 3 mice per group). **(H)** Representative traces showing repetitive firing of APs after prolonged (5s) current injection (2x Rheobase). **(I)** CGRP release from cultured DRG neurons induced by 15 min depolarization with KCl (10, or 40 mM) or capsaicin (10 µM, Cap) measured by EIA assay. KCl-depolarization was performed also in the presence BayK (1 µM) nimodipine (10 µM, Nimo). **(J-K)** HC imaging-based quantification of CGRP expression in overnight cultures of DRG neurons from Cacna1c^fl/fl^ and SNS-Cacna1c^-/-^ mice. Electrophysiological data in (B) and (D-H) are medians with 95% percentiles; Mann-Whitney U test. CGRP EIA data in (I) are means ± SEM; n = 4, two-way ANOVA with Bonferroni’s test. HCI data in (J) are means ± SEM; n = 4; Welch’s t-test. \**p* < 0.05; \*\**p* < 0.01; \*\*\**p* < 0.001 indicate significance levels between the indicated conditions.

Next, we quantified basal electrophysiological properties of Cav1.2-deficient neurons. Neither the rheobase nor basic characteristics of action potentials such as the overshoot or duration of repolarization were significantly altered in the tested neurons of SNS-Cacna1c^-/-^ mice compared to controls (**Figures 7C-F**). However, the Cav1.2-deficient neurons showed substantially impaired repetitive firing after prolonged current injection (**Figures 7G-H**). In total, 16 of 18 wildtype neurons, but only 7 of 22 Cav1.2-deficient neurons, showed repetitive firing during a 5 sec test period. In conclusion, Cav1.2-deficiency severely impairs the repetitive firing upon neuronal excitation.

### Calcium influx through Ca_V_1.2 leads to CGRP release

Calcium influx during depolarization induces the release of neuropeptides such as calcitonin gene-related peptide (CGRP) from peripheral and central terminals of sensory neurons by classical calcium-dependent exocytosis ^94^. In the periphery, CGRP acts as a potent vasodilator during neurogenic inflammation ^95^. At central synapse in the spinal cord, CGRP acts as an excitatory neurotransmitter and its release is mainly attributed to the N-type calcium channels (Cav2.2) enriched at presynaptic nerve terminals ^72,96,97^, which are the targets of ω-conotoxin based analgesics such as Ziconotide ^98^. However, CGRP can also be released at peripheral terminals, along neurites and even cell bodies, where Cav1 channels are expressed at high levels ^99^. In SNS-Cacna1c^-/-^ mice, our transcriptome analysis of DRGs revealed downregulation not only of Cav1.2 but also of Cav2.2, along with proteins associated with calcium-dependent vesicle secretion and recycling. We therefore assessed the impact of Cav1.2-deficiency on CGRP release from DRG neurons using a conventional enzyme immunoassay.

Indeed, KCl-depolarization for 15 min led to dose-dependent CGRP release from DRG neurons of both genotypes that significantly exceeded the capsaicin-induced CGRP release (**Figure 7I**). However, CGRP release was already evident at mild KCl-depolarization (10 mM KCl) in control DRG neurons, but not Cav1.2-deficient DRG neurons. In addition, the amplitude of CGRP release was lower in Cav1.2-deficient DRG neurons after robust KCl-depolarization (40 mM KCl). The Cav1 agonist BayK reinforced the CGRP release in control neurons after mild KCl-depolarization, but not in Cav1.2-deficient DRG neurons. Supporting this, also the Cav1 antagonist nimodipine significantly inhibited the depolarization-dependent CGRP release in control neurons only indicating that calcium influx through Cav1.2 is sufficient to induce CGRP release from cultured DRG neurons.

### SNS-Cacna1c^-/-^ mice show increased heat nociception

Next, we assessed the role of Cav1.2-deficiency for pain processing *in vivo*. For this, we compared the behavior of SNS-Cacna1c^-/-^ mice with littermate Cacna1c^fl/fl^ control mice in several pain models. Motor coordination was not altered in SNS-Cacna1c^-/-^ mice in an accelerating rotarod test: SNS-Cacna1c^-/-^ mice, 276 s; control mice, 253 s; *P* = 0.17; *n* = 10; **Figure 8A**), allowing the analysis of their immediate responses to acute noxious stimuli. In the hot plate test, in which the animals are placed on a heated surface followed by measuring the time until the mouse shows nocifensive behavior, SNS-Cacna1c^-/-^ mice had significantly lower response latencies compared to control mice when the plate was set to 52°C or 54°C, indicating increased heat nociception (**Figure 8B**). Similar results were obtained by analyzing an independent cohort of mice in a different laboratory (**Figure S9A**). However, the withdrawal latencies after stimulating the tail with a mild or high intensity beam were not altered in the SNS-Cacna1c^-/-^ mice as compared to control mice (**Figure 8C**). On a cold plate set to 5°C or 10°C, the time spent lifting one or both forepaws in 60 s was similar between genotypes (**Figure 8D**). Furthermore, the mechanical pain threshold, as assessed by stimulation of the hindpaws with von Frey filaments, was normal in SNS-Cacna1c^-/-^ mice (**Figure 8E**). Measuring the mechanical sensitivity by the Pincher test on the mouse tail also did not reveal alterations concerning the sensation of mechanical stimuli (**Figure 8F**). Together, these data suggest that the sensitivity to noxious heat is increased in SNS-Cacna1c ^-/-^ mice, while cold sensation and mechanosensation are normal.

**Figure 8.**
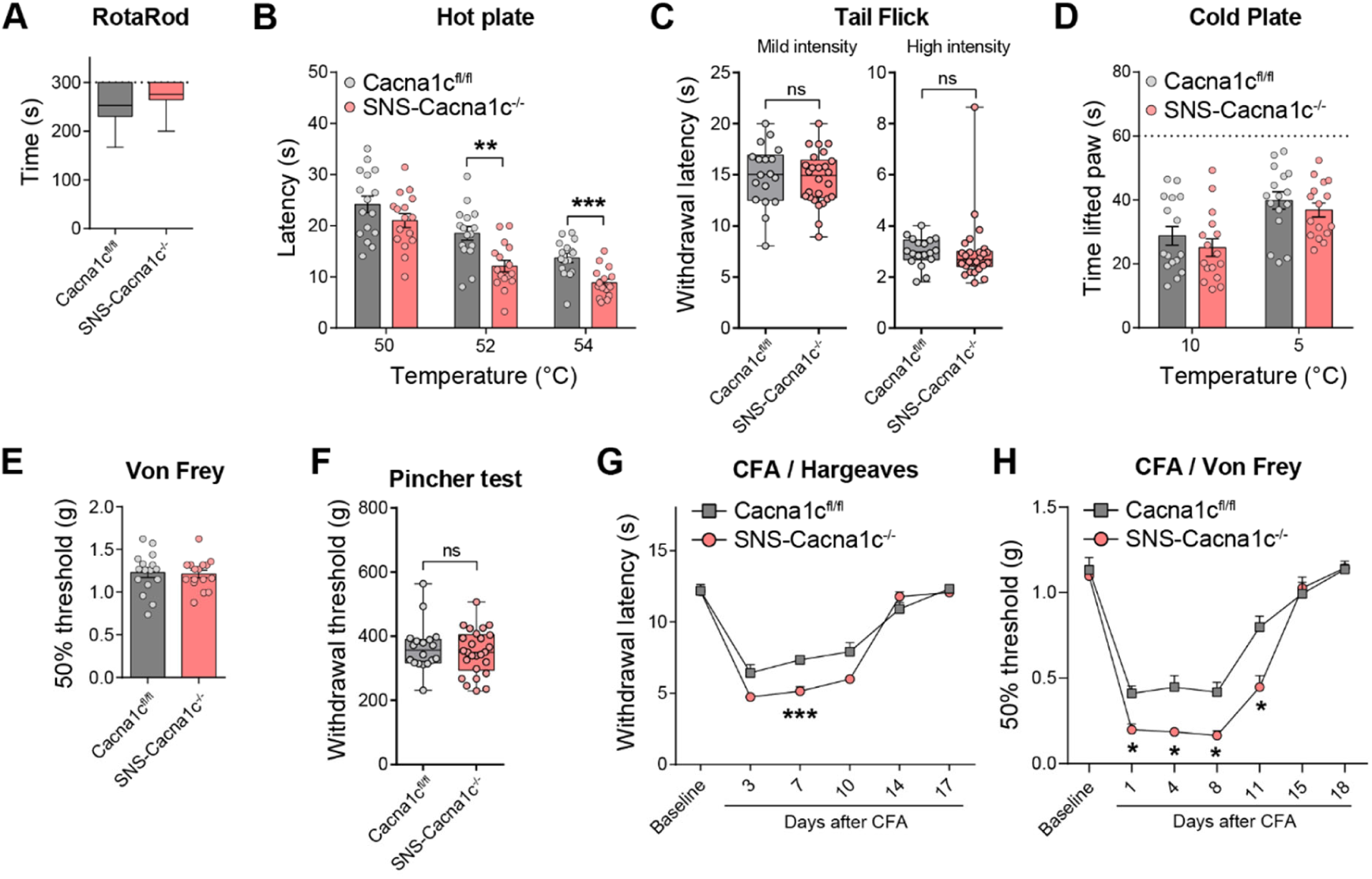
SNS-Cacna1c^-/-^ mice show increased heat nociception and CFA-induced inflammatory pain. **(A)** Fall-off latencies of SNS-Cacna1c^-/-^ and Cacna1c^fl/fl^ control mice in an accelerating rotarod test. The dotted line indicates the cutoff time (n = 10, male and female mice, Mann Whitney test). **(B)** Withdrawal latencies showing a nociceptive response on a hot plate set to 50°C, 52°C, or 54°C (n = 16, male and female mice, Multiple t-test, Holm-Sidak method). **(C)** Thermal heat sensitivity measured by Tail Flick test on the mice tail with a mild (left panel) or high heat intensity beam (right panel) (n = 18 Cacna1c^fl/fl^ and 26 SNS-Cacna1c^-/-^, male and female mice, unpaired t-tests with Welch correction). **(D)** Total time of forepaw lifts from a cold plate set to 10°C or 5°C within an observation time of 60 s (n = 16, male and female mice, Multiple t-test, Holm-Sidak method). **(E)** Mechanical pain threshold determined using von Frey filaments (n = 16, male and female mice, t-test). **(F)** Mechanical sensitivity measured by Pincher test on the mouse tail (n = 18 Cacna1c^fl/fl^ and 26 SNS-Cacna1c^-/-^, male and female mice, unpaired t-tests with Welch correction). **(G)** Thermal hypersensitivity assessed by radiant heat stimulation (Hargreaves method) after injection of complete Freund’s adjuvant (CFA) into a hindpaw (n = 8, male and female mice, repeated measures two-way ANOVA with Sidak post hoc test). **(H)** Mechanical hypersensitivity measured using von Frey filaments after injection of CFA into a hindpaw. Data in (A), (C), and (F) are presented as median with interquartile range, other data are presented as mean ± SEM. \**p* < 0.05; \*\**p* < 0.01; \*\*\**p* < 0.001 indicate significance levels between the indicated groups.

### Cav1.2-deficiency in nociceptors exacerbates inflammatory but not neuropathic pain

To model persistent inflammatory pain, we injected Complete Freund’s Adjuvant (CFA) into a hindpaw and analyzed the extent of thermal and mechanical hypersensitivity. Of note, thermal hypersensitivity measured using the Hargreaves test was significantly increased in SNS-Cacna1c^-/-^ mice compared to control littermates at days 3-10 after the CFA injection (**Figure 8G**). In addition, the mechanical pain threshold determined by von Frey filaments was decreased at days 1-11 after CFA injection indicating increased CFA-induced mechanical hyperalgesia in SNS-Cacna1c^-/-^ mice (**Figure 8F**).

To assess whether Cav1.2-deficiency also affects the development of neuropathic pain, we applied the spared nerve injury (SNI) model. In this model, the common peroneal and tibial nerves are injured, producing mechanical allodynia in the skin territory of the spared, intact sural nerve ^100^. The SNI procedure led to a consistent increase in mechanical, heat, and cold sensitivity in SNS-Cacna1c^-/-^ mice and floxed control littermates. However, the mechanical and thermal hypersensitivity was similar in both genotypes (**Figures S10B-D**). These *in vivo* experiments revealed that *Cacna1c* expressed in nociceptors contributes to hypersensitivity during persistent inflammatory but not neuropathic pain.

## Discussion

Within this study, we investigated the role of Cav1.2 in peripheral nociceptors to determine its contribution to pain perception. Specifically, our goal was to elucidate whether Cav1-dependent signaling induces E-T coupling in nociceptors and thereby influences the development of chronic inflammatory or neuropathic pain, a context in which E-T coupling remains unexplored. To accomplish this, we made use of a conditional mouse model that lacks Cav1.2 in Nav1.8-expressing nociceptive sensory neurons and performed functional studies in isolated sensory neurons *ex vivo* and pain models *in vivo*.

We found that KCl-depolarization of isolated DRG neurons induces CREB activity in a Cav1, PKA, and CaN-dependent manner indicating that basic mechanisms mediating E-T coupling act also in nociceptors ^6^. Genetic or pharmacological inhibition of Cav1.2 reduced calcium influx, calcium-dependent kinase signaling, and CREB phosphorylation after KCl-depolarization. However, Cav1.2-deficiency reduced the calcium influx only in the presence of lower KCl concentrations, suggesting that calcium influx via Cav1.2 occurs already during mild KCl-depolarization, and is masked by calcium influx through other VGCCs during stronger KCl-depolarization. Intriguingly, blocking Cav2 channels with aga- and conotoxins alone did not reduce the calcium influx. Instead, combining all Cav2 blockers with nimodipine was necessary for substantial inhibition of calcium influx indicating that the total amount of calcium in- and efflux is tightly controlled.

In the same line of evidence, Cav1.2-deficiency or pharmacological inhibition produced a rightward shift of the PKA-II or pCREB dose-response relationship, which was more pronounced in the presence of the Cav1 agonist BayK. Again, inhibitors of Cav2 channels alone were ineffective and combination with nimodipine was required to induce a synergistic inhibitory effect. This demonstrates that calcium influx via alternate VGCCs, followed by diffusion into Cav1 nanodomains, can activate Cav1-dependent signing and masks genotype-associated differences concerning PKA-II and CREB activation during strong KCl-depolarization. Similar observations have already been made in superior cervical ganglion (SCG) neurons, in which mild KCl-depolarization signal exclusively through Cav1 to CaMKII and CREB, whereas stronger KCl-depolarization recruits Cav2 channels ^9,10,23^.

Concerning the mechanism of Cav1.2-dependent E-T coupling, previous studies in other neuronal cell models indicate that Ca^2+^ influx into Cav1.2 nanodomains is required ^9^, but a voltage-dependent conformational change (VΔC) provides a necessary additional signal ^25^. Our data is in accord with this model, but further studies are needed to claim that a VΔC is necessary for effective E-T coupling in nociceptors. Further, research on E-T coupling in sympathetic and CNS neurons established a central role of CaMKII and IV ^9,24,36,39^. Formation of CaMKII puncta near the cell surface suggested that calcium influx is required for mobilization of CaMKII from F-actin followed by VΔC-dependent accumulation at Cav1.2 ^25^. CaMKII is highly expressed in a subset of DRG neurons ^101^ and studied in the context of central sensitization and LTP ^14,102^. However, CaMKII or IV inhibitors did not inhibit KCl-induced pCREB, even if we tested long-term cultures of DRG neurons from newborn rats in an attempt to resemble the culture conditions used in the studies of others. Therefore, methods must be established to assess the role of CaMK activity for E-T coupling in DRG neurons in the future.

Despite the molecular details mediating E-T coupling, we demonstrate that modulation of Cav1 activity leads to transcriptional regulation in cultured DGR neurons *in vitro* and whole DRGs *in vivo*. Many of the genes identified after *in vitro* stimulation of DRG neurons have been also detected in other neuronal cell models after acute depolarization ^51,56^. The expression of these IEGs does not depend on *de novo* synthesis of other transcriptional regulators and is considered as a marker signature to track neuronal activity, for instance in the context of memory formation ^103^. Strikingly, most of the activity-regulated transcripts showed opposing regulation in the presence of Cav1 agonist or antagonist, respectively.

A network analysis of the Cav1-associated transcriptomic changes provides strong evidence for a specific modulation of transcription by Cav1-dependent signaling. Indeed, for several of the identified proteins and interactions, a role in nociception and different forms of pain has been previously identified. However, regulation of IEGs represents a first wave of activity-dependent transcriptional changes initiating long-term changes to establish an altered homeostatic state. Therefore, it is difficult to assess the net impact of these changes on the activity profiles of the affected nociceptors. In an attempt to identify changes associated with long-term Cav1.2-dependent E-T coupling, we analyzed the transcriptome of whole DRGs of SNS-Cacna1c^-/-^ mice and control littermates by bulk RNA-Seq. Using HCI, we first verified that these mice do not show substantial alterations regarding the composition of neuronal subgroups within the DRG. Although the of neuronal subgroups was unchanged, > 500 genes showed differential expression at a 1.5-fold cut-off. This finding strongly supports that Cav1.2-dependent E-T coupling has an important function for the homeostatic level of gene expression in sensory DRG neurons. However, due to the different time scale, we did not detect a substantial overlap of genes identified after acute depolarization *in vitro* and long-term knockout *in vivo*. Further experiments need to address this by more in-depth kinetic experiments with multiple points, including longer time points.

Unexpectedly, deletion of Cav1.2 did not induce compensatory upregulation of Cav1.3 in sensory neurons. This was observed in cardiomyocytes ^91^, but not after conditional deletion of Cav1.2 in CNS neurons ^104^. Of note, many of the identified genes contribute to the generation of action potentials and the synaptic release of neuropeptides. The identified transcriptional changes therefore potentially indicate that Cav1.2-dependent E-T coupling represents a mechanism that regulates the homeostatic excitability of the neuron. To test for changes in the excitability, we performed patch-clamp recordings. Unexpectedly, this analysis revealed a highly significant reduction of the repetitive AP firing in DRG neurons from SNS-Cacna1c^-/-^ mice whereas basic parameters were not affected. The mechanism how neurons regulate their spiking (firing) frequency is complex and depends on the interplay between several ion channels that dictate the refractory period, afterhyperpolarizations (AHPs), and afterdepolarizations (ADPs) that followed each action potential ^105^. Taken that the deletion Cav1.2 was associated with a dysregulation of several ion channels, it is difficult to speculate which of the changes in SNS-Cacna1c^-/-^ mice are causal for this phenotype. In additon, we have recently shown that Cav1.2-dependent calcium influx activates PKA-II in DRG neurons ^5^; a finding that we further investigated within this study. PKA is considered as one of the central kinases modulating the activity of various sodium, calcium, and potassium channels. Further studies need to address whether the lack of PKA-dependent ion channel phosphorylation underlies the impaired capability of repetitive firing in Cav1.2-deficient neurons.

As the second integrative readout, we tested whether Cav1.2-deficiency modulates excitation-secretion (E-S) coupling in cultured DRG neurons. So far, this is mainly attributed to N-type channels (Cav2.2) enriched at presynaptic nerve terminals ^72^, since genetic deletion ^96,97^ or spinal inhibition with ω-conotoxins has substantial analgesic effects ^106–110^. However, in cultured DRG neurons of SNS-Cacna1c^-/-^ mice, we observed a substantial impairment of depolarization-dependent CGRP release. The difference between genotypes directly correlated with Cav1.2 activity, as it was increased by Cav1 agonist and diminished by Cav1 antagonist, respectively. Further studies are required to elucidate the in vivo relevance of Cav1-dependent CGRP release; however, a cooperative interaction between Cav1 and Cav2 channels may be necessary.

Our findings add to the increasing knowledge that VGCC isoforms have distinct but partially overlapping functionalities during nociception. To our surprise, and not expected from the *in vitro* data showing diminished calcium influx, kinase activity, repetitive AP firing, and CGRP release, we observed increased basal heat nociception as well as CFA-induced thermal hypersensitivity and mechanical allodynia in SNS-Cacna1c^-/-^ mice *in vivo*. The increased nociception in SNS-Cacna1c^-/-^ mice is potentially attributed to the lack of transcriptional changes induced by Cav1.2-dependent E-T coupling in wildtype mice. Therefore, Cav1.2-dependent E-T coupling may counteract nociception *in vivo*. For instance, downregulation of multiple potassium channels such as *Kcnb1, Kcng3, Kcnh6, Kcnk18* and *Kcnt1* could well explain the overall increase in heat and inflammatory pain perception in the SNS-Cacna1c^-/-^ mice. However, while long-term genetic Cav1.2 deficiency exacerbates pain perception, Cav1 antagonists exhibit acute analgesic effects. For instance, local injection of KCl into the hindpaw of rats induces mechanical hyperalgesia, which can be alleviated by the Cav1 antagonist verapamil ^5^. In addition, a broad-spectrum dihydropyridine inhibitor of VGCCs has analgesic effects partly due to Cav1 inhibition ^111^. Our findings resemble a study investigating fear memory formation. Acute pharmacological block of Cav1.2, but not genetic inactivation in CNS neurons, impaired the acquisition of conditioned fear memory in mice ^104^. There, Cav1.2-deficiency changed the molecular mechanism of Cav1.2-dependent Hebbian LTP in the lateral amygdala by upregulation of GluA1-containing calcium-permeable AMPARs ^104^. Calcium influx via Cav1.2 therefore leads to homeostatic changes in postsynaptic plasticity. Here we demonstrate that loss of Cav1.2 also induces homeostatic changes in presynaptic sensory neurons. Further studies should now clarify whether the activation or inhibition of Cav1.2-dependent E-T coupling has therapeutic benefit in certain pain conditions.

## Resource Availability

### Lead contact

Further information and requests for resources and reagents should be directed to and will be fulfilled by the lead contacts, Joerg Isensee and Tim Hucho.

## Material availability

This study did not generate new, unique reagents.

## Data and code availability

RNA sequencing data are publicly available at the EMBL-EBI repository ArrayExpress with the accession numbers E-MTAB-15261 and E-MTAB-15237. This paper does not report original code. Any additional information required to reanalyze the data reported in this paper is available from the lead contact upon request.

## Acknowledgements

JI, PNO, and TH were supported by grants from the German Research Foundation (DFG 516750869). ICAPS was supported by the PRACTIS Clinician Scientist Program, funded by the Hannover Medical School and the German Research Foundation (DFG ME 3696/3)

## Author contributions

Conceptualization: JI, TH Methodology: JI, MS

Formal analysis: JI, CWL, BD, FB, PFB

Investigation: JI, PE, PC, ML, MS, PNO, RD, ME, IS, RL

Resources: FB, PFB

Data Curation: JI, CWL, BD, FB, RL, PFB Writing - Original Draft: JI, PNO, AS, PFB, TH

Writing - Review & Editing: JI, PNO, PFB, AS, AL, TH Supervision: MRS, PFB, AL, AS, TH

Funding acquisition: JI, PFB, TH

## Data and materials availability

For material and correspondence please contact Jörg Isensee, Tim Hucho.

## Materials and Methods

### Key resources table

**Table 1.**
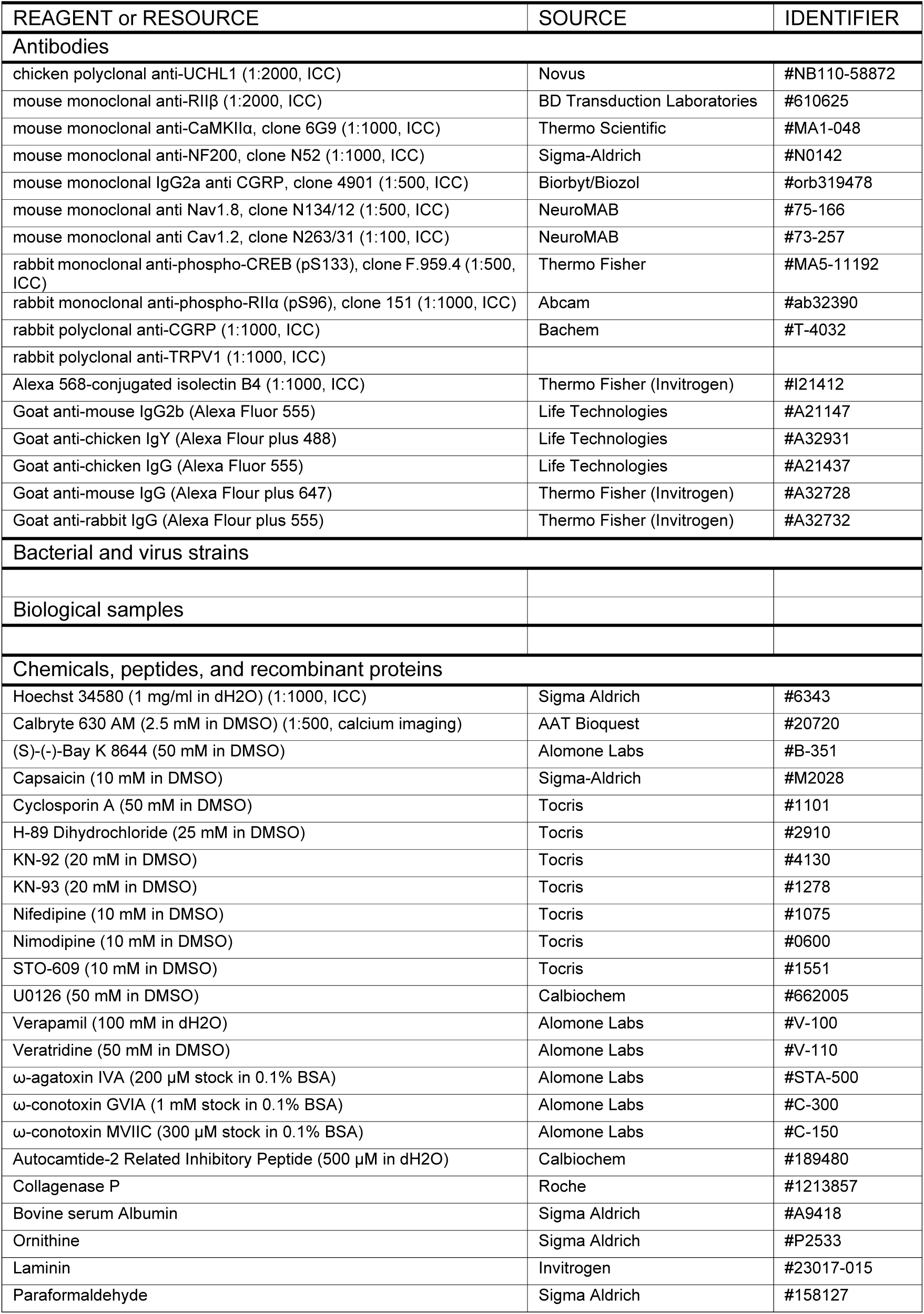
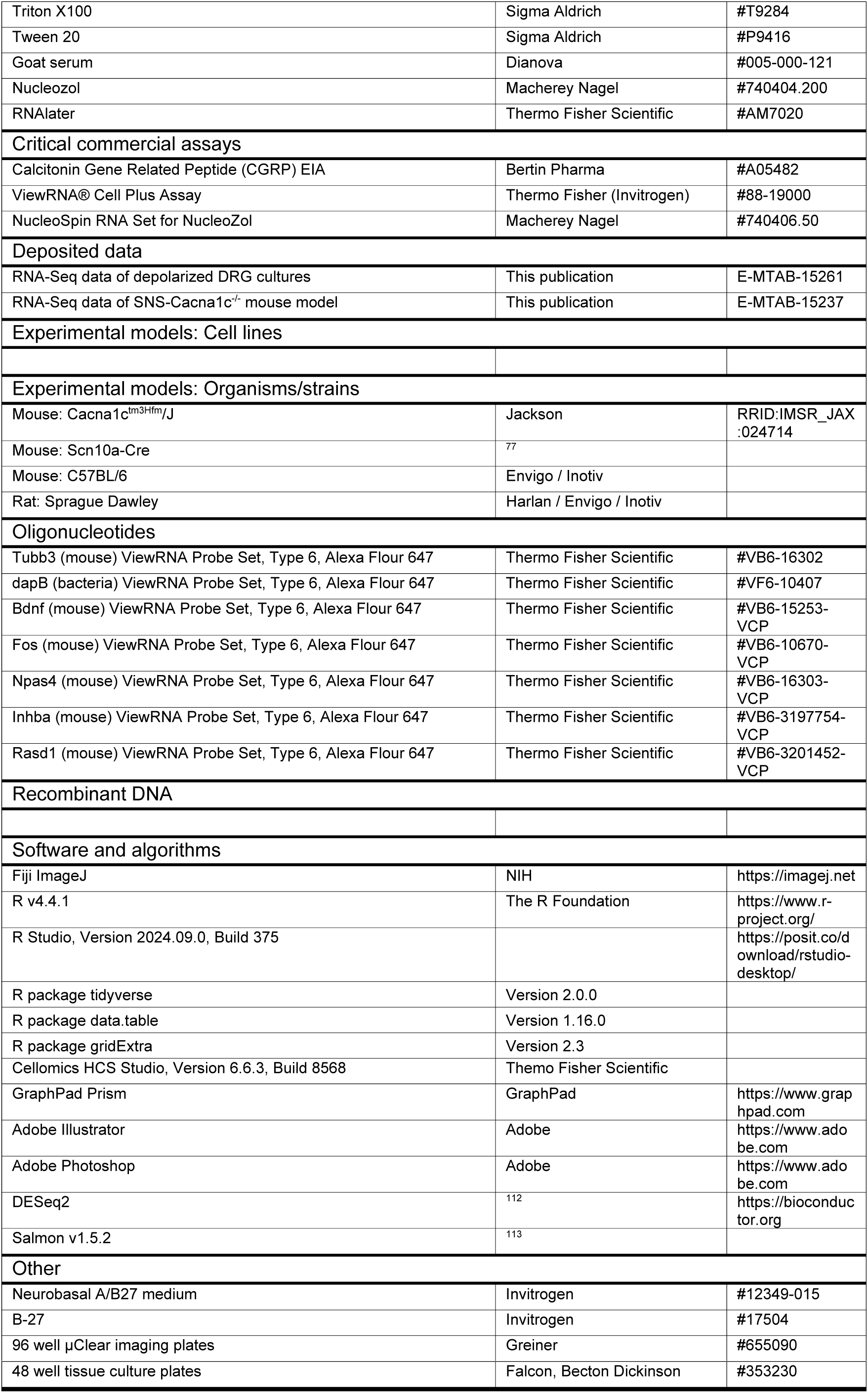
Demographic data comparing those who tested positive for anti-LID-1 antibody and those who tested negative.

### Experimental model and study participant details

Male Sprague Dawley rats (200-225 g, aged 8-10 weeks) were obtained from Harlan, Envigo or Inotiv. Rats were kept in a temperature- and humidity-controlled animal care facility at the University Hospital of Cologne on a 12-h light/dark cycle and provided with food and water *ad libitum*. Tissue-specific conditional Cav1.2 mutant mice were generated by crossing floxed mice, which carry loxP sites flanking exons 14-15 of the murine *Cacna1c* gene ^76^ (Cacna1c^tm3Hfm^, JAX No. 024714), with mice carrying a sensory neuron-specific (SNS) Cre recombinase under control of the *Scn10a* (Nav1.8) promoter^77^. Offspring were kept on a 12 h light/dark cycle and provided with food and water *ad libitum*. For cellular experiments, 8-24 weeks old female and male mice were sacrificed between 9-12 a.m. by CO_2_ intoxication. Dorsal root ganglia (cervical, lumbar and thoracic) were removed within 30 min per animal. Nociceptive testing was performed using 10-16 weeks old female and male mice. All experiments adhered to the guidelines of the International Association for the Study of Pain and to the ARRIVE (Animal Research: Reporting on In Vivo Experiments) guidelines, and were approved by the local Ethics Committee for Animal Research (Regierungspräsidium Darmstadt, Germany; Landesamt für Natur, Umwelt und Verbraucherschutz Nordrhein-Westfalen, Germany; Tissue removal (DRG) from adult mice was approved by the Animal Protection Officer of Hannover Medical School [§4-Anzeige, Nummer 3741, Approval date 1 January 2024]; Committee on Animal Experimentation of the Canton de Vaud, Switzerland, approved animal protocol number 34684/VD1339.10d). Effort was made to minimize the number of animals used and their suffering.

### Antibodies

The following antibodies were used in this study: chicken polyclonal antibodies anti-UCHL1 (1:2000, Novus, #NB110-58872), rabbit monoclonal antibody anti-phospho-CREB pS133 (1:500, clone F.959.4, Thermo Fisher, #MA5-11192), rabbit monoclonal antibody anti-phospho-RIIα (pS96) (1:1000, clone 151, Abcam, #ab32390), mouse monoclonal antibody anti-RIIβ (1:2000, BD Transduction Laboratories, #610625), mouse monoclonal anti-CaMKIIα (1:1000, Thermo Scientific, clone 6G9, #MA1-048), rabbit anti-CGRP antibody (1:1000, Bachem, #T-4032), mouse monoclonal anti-NF200 (1:1000, Sigma-Aldrich, clone N52, #N0142), mouse monoclonal IgG2a anti CGRP (1:500, Biorbyt/ Biozol, clone 4901, #orb319478), mouse monoclonal anti Nav1.8 (1:500, NeuroMAB, clone N134/12 75-166), mo-Cav1.2_CloneN263/31: mouse monoclonal anti Cav1.2 (1:200, NeuroMAB, clone N263/31, #73-257), rabbit polyclonal anti-TRPV1 (1:1000, Alomone, #ACC-030), and highly cross-adsorbed Alexa 647, 594, and 488 conjugated secondary antibodies (Invitrogen, Carlsbad, CA). IB4 was detected using Alexa 568-conjugated isolectin B4 (1:1000, Invitrogen, #I21412).

### Reagents

Stock solutions were prepared as indicated in parentheses of the following compounds: Autocamtide-2 Related Inhibitory Peptide (AIP, Calbiochem, #189480, 500 µM in dH2O), (S)-(-)-Bay K 8644 (BayK, Alomone Labs, #B-351, 50 mM in DMSO), Capsaicin (Cap, Sigma-Aldrich, #M2028, 10 mM in DMSO), Cyclosporin A (CsA, Tocris, #1101, 50 mM in DMSO), H-89 Dihydrochloride (H89, Tocris, #2910, 25 mM in DMSO), KN93 (Tocris, #1278; 20 mM in DMSO), Nifedipine (Nife, Tocris, #1075, 10 mM in DMSO), Nimodipine (Nimo, Tocris, #0600, 10 mM in DMSO), STO-609 (Tocris, #1551, 10 mM in DMSO), U0126 (Calbiochem, #662005, 50 mM in DMSO), Verapamil (VP, Alomone Labs, #V-100, 100 mM stock in dH2O), Veratridine (VT, Alomone Labs; #V-110; 50 mM stock in DMSO), ω-agatoxin IVA (Alomone Labs, #STA-500, 200 µM stock in 0.1% BSA), ω-conotoxin GVIA (Alomone Labs, # C-300, 1 mM stock in 0.1% BSA), ω-conotoxin MVIIC (Alomone Labs, #C-150, 300 µM stock in 0.1% BSA).

### Dorsal root ganglion neuron cultures

DRGs were isolated, pooled, and incubated in Neurobasal A/B27 medium (Invitrogen, #12349-015) containing collagenase P (Roche, #1213857) (0.2 U/ml, 1 h, 37 °C, 5% CO_2_). The dorsal root ganglia were dissociated by trituration with fire-polished Pasteur pipettes. Axon stumps and disrupted cells were removed by bovine serum albumin (BSA) gradient centrifugation (15% BSA, 120 *g*, 8 min). Viable cells were resuspended in Neurobasal A/B27 medium, plated in poly-L-ornithine (0.1 mg/ml)/laminin (5 µg/ml)-precoated 96-well imaging plates (Greiner, #655090) and incubated overnight (37°C, 5% CO_2_). Neuron density was 1500 neurons/cm^2^.

### Stimulation of dorsal root ganglion neurons

DRG neurons were stimulated 24 h after isolation in 96-well imaging plates. Compounds were dissolved in 12.5 µl PBS, pH 7.4, in 96-well V-bottom plates, mixed with 50 µl medium from the culture wells, and added back to the same wells. Stimulations were performed with automated 8 channel pipettes (Eppendorf) at low dispense speed on heated blocks, stimulated cells were placed back in the incubator. Kinetic experiments were performed in the reverse order starting with the longest time point. Cav1 inhibitors or agonists such as verapamil, nimodipine, or (S)-(-)-Bay K 8644 that reach their target sites from the extracellular space were preincubated for 10 min before depolarization. Inhibitors of intracellular proteins were preincubated for 30 min to 1 h before depolarization. The stimulated cells were fixed for 10 min at room temperature (RT) by adding 100 µl 8% paraformaldehyde resulting in a final concentration of 4%.

### Immunocytochemistry

Fixed cells were treated with goat serum blocking solution (2% goat serum, 1% BSA, 0.1% Triton X-100, 0.05% Tween 20, 1 h, RT) and incubated with respective primary antibodies diluted in 1% BSA in PBS at 4°C overnight. After three washes with PBS (30 min, RT), cells were incubated with secondary Alexa dye-coupled antibodies (1:1000, 1h, RT). After three final washes (30 min, RT), wells of 96-well plates were filled with PBS, sealed, and stored at 4°C until scanning.

### Single molecule Fluorescence In Situ Hybridization (smFISH)

We applied the ViewRNA Cell Plus Assay (Thermo Fisher Scientific, #88-19000) to combine the immunocytochemical detection of the neuronal marker UCHL1 and fluorescent *in situ* hybridization against respective target transcripts *Bdnf*, *Fos*, *Npas4*, *Inhba*, and *Rasd1*. In brief, mouse DRG cultures were prepared in poly-L-ornithine (0.1 mg/ml)/laminin (5 µg/ml)-precoated 96-well imaging plates (Greiner, #655090) and incubated overnight (37 °C, 5% CO_2_). The cultures were stimulated with 40 mM KCl as described above. The staining procedure was performed according to the manufacturer’s instructions. The following Type 6 ViewRNA probe sets were used: Tubb3 (mouse, VB6-16302-VCP), dapB (bacteria, VF6-10407-VCP), Bdnf (mouse, VB6-15253-VCP), Fos (mouse, VB6-10670-VCP), Npas4 (mouse, VB6-16303-VCP), Inhba (mouse, VB6-3197754-VCP), Rasd1 (mouse, VB6-3201452-VCP).

### High content imaging

We used a Cellomics CX7 LZR microscope equipped with a 16-bit Photometrics camera and laser light source to scan stained cultures of DRG neurons in 96-well imaging plates. 2x2 binned images (1104 x 1104 pixels) were acquired with a 10× (NA = 0.4) X-line objective (Olympus) and analyzed using the Cellomics software package. Briefly, images of UCHL1 staining were background corrected (low pass filtration), converted to binary image masks (fixed threshold), segmented (geometric method), and neurons were identified by the object selection parameters: size of 80-7500 µm^2^, circularity (perimeter^2^ / 4π area) of 1-3; length-to-width ratio of 1-2; average intensity of 800-12000, and total intensity of 2x10^5^-5x10^7^. The image masks were then used to quantify signal in other channels. To calculate spill-over between fluorescence channels, three respective controls were prepared for each triple staining: (i) UCHL1 alone, (ii) UCHL1 + antibody 1, and (iii) UCHL1 + antibody 2. Raw fluorescence data of the controls were used to calculate the slope of best fit straight lines by linear regression, which was then used to compensate spill-over as described previously ^114^. Compensated data were scaled to a mean value of 1 for the unstimulated cells to adjust for variability between experimental days. One and two-dimensional probability density plots were generated using R packages ^115^. Gating of subpopulations was performed by setting thresholds at local minima of probability density plots.

### Calcium imaging

DRG neurons were seeded in poly-L-ornithine (0.1 mg/ml)/laminin (5 µg/ml)-precoated 96-well µclear imaging plates (Greiner, #655090) in 100 µl supplemented NB medium and cultured overnight. The cells were loaded for 60 min with 5 µM Calbryte 630 AM diluted in medium. The cells were carefully washed twice with 37 C warm saline solution (140 mM sodium chloride, 5 mM potassium chloride, 2 mM calcium chloride, 1 mM magnesium chloride, 10 mM D-glucose, 10 mM HEPES, pH7.4 with sodium hydroxide). Images were acquired at 3 Hz with a Leica DMI6000BSI microscope using the 5x objective. After a baseline scan (20 images), the stimulus was applied 5-fold concentrated in 50 µl volume with a gel loading tip. After acquiring 120 images, KCl (40 mM final conc.) was applied to depolarize the neurons (60 images). The images were analyzed using a custom Image J script after background correction (rolling ball). The images 160-200 were auto thresholded (Li dark algorithm) and used to generate an image mask with the particle analyzer. The image mask was then applied to quantify the fluorescence intensities in all objects and time points. Raw fluorescence intensities were normalized to the mean of the first 10 images and ΔF/F values were calculated. Only neurons showing a response to the compound and/or KCl were considered for further data processing.

### CGRP release assay

DRG neurons were plated in 48-well plates (Falcon, Becton Dickinson, #353230) pre-coated with poly-L-ornithine (0.1 mg/ml) and laminin (5 µg/ml) in 250 µl of supplemented Neurobasal (NB) medium and cultured for 2 days. The culture medium was refreshed four times with 100 µl of fresh medium per exchange. Following a 30 min incubation at 37°C, 100 µl of medium was removed from each well, resulting in a final volume of 150 µl. Cells were pre-incubated for 15 min with (S)-(-)-Bay K 8644, nimodipine, or DMSO (vehicle control) by adding 50 µl of a 4x concentrated solution prepared in fresh medium. Subsequently, stimulation was performed for 15 min by adding 50 µl of a 5x concentrated stimulus (KCl or capsaicin, diluted in medium). The supernatants were collected immediately and placed on ice. Samples were centrifuged for 1 min at 1000 g at 4°C and stored at -80°C. Calcitonin Gene-Related Peptide (CGRP) concentrations were determined in duplicates using a commercial enzyme immunoassay (EIA) kit (Rat CGRP EIA, Bertin Pharma, Cat# A05482) according to the manufacturer’s protocol.

### Behavioral testing

All behavioral studies were conducted in littermate mice and by an observer blinded for the genotype and treatment of the animals. Prior to the experiments mice were acclimated to the testing room for at least 1 h.

#### Accelerating rotarod test

Motor coordination was assessed with a rotarod treadmill for mice (Ugo Basile, Gemonio, Italy) at increasing rotating speed from 4-40 rpm over 300 s, with a cutoff time at 300 s. All mice had training sessions for consecutive days before the experiment. The mean of three trials was used for evaluation.

#### Hot plate test

Mice were placed on a heated metal surface (hot/cold plate, Ugo Basile) and the time between placement and a nociceptive behavior (licking or shaking of a hindpaw, jumping) was recorded. Temperatures of 50°C, 52°C, and 54°C were applied with cutoff times of 60 s, 40 s and 20 s, respectively. Only one test per animal was performed.

#### Cold plate test

Mice were placed on a cold plate (Ugo Basile) maintained at 5 °C or 10°C and the total time that the animal spent lifting one or both forepaws during 60 s after placement was measured ^116^. One test per animal was performed.

#### Von Frey filament test

For determination of mechanical thresholds, animals were placed in a chamber with a wire mesh floor and allowed to habituate for at least 30 minutes before testing. Calibrated von Frey filaments (Ugo Basile) ranging from 0.4 to 19.6 mN (0.04–2.0 g) were applied to the plantar surface of a hindpaw, and measurements were started with the 0.4-g filament. A clear paw withdrawal, shaking, or licking during or immediately after the stimulus (up to 3 seconds after the filament was bowed) was defined as a response. The 50% withdrawal thresholds were calculated using the online tool, “Up-down method for von Frey experiments” (https://bioapps.shinyapps.io/von_frey_app/) ^117^.

#### Paw licking and hypersensitivity induced by capsaicin and AITC

Baseline measurements using the von Frey filament test were performed as described above. Then animals were habituated in a plexiglass cylinder for at least 30 min. Capsaicin (Sigma Aldrich; 5 μg ^118^ or allyl isothiocyanate (AITC; Sigma Aldrich; 10 mM), dissolved in 20 μl PBS containing 2% dimethyl sulfoxide, were injected into the plantar surface of a hindpaw. Immediately thereafter animals were placed in the plexiglass cylinder and the time spent licking the injected paw was recorded for 20 min (capsaicin) or 30 min (AITC) using a stop watch. Then animals were placed to the chamber with a wire mesh floor and the von Frey filament test was performed as described above over 72 h.

#### CFA-induced paw inflammation

Twenty microliter of complete Freund’s adjuvant (CFA; containing 1 mg/ml heat-killed *Mycobacterium tuberculosis* in paraffin oil 85% and mannide monooleate 15%; Sigma-Aldrich) was subcutaneously injected into the plantar side of a hindpaw ^119^. Sensitivity to thermal stimulation ^120^ was measured using a using a Plantar Test Instrument (IITC). Animals were placed in observation chambers on top of a temperature controlled, warm glass surface. The radiant heat source was aimed at the plantar surface of the mid-hindpaw of an inactive mouse. The paw withdrawal latency was calculated as the mean of 5 consecutive trials with at least 30 s in-between. Mechanical sensitivity of the hindpaw was determined using the von Frey filament test as described above.

#### Pincher test

The Pincher test consists of a pair of blunt forceps (15 cm long; flat contact area: 7 mm x 1.5 mm with smooth edges) equipped with two strain gauges connected to a modified electronic dynamometer (Bioseb). The mice are gently restrained in a conical plastic cloth. The tips of the forceps are placed around the tail of the tested mice, and the force applied was incremented by hand until a withdrawal response occurred (limit max 15 g). The measurement is repeated 3 times, and the mean force (in grams) inducing withdrawal is calculated. The testing last for 5 min.

#### Tail Flick assay

The Tail-flick assay is conducted using a tail-flick analgesia meter and the mice are gently restrained in a conical plastic cloth. The latency of response (in seconds) is recorded at two different light-beam intensities (Mild:4 and High:7, on two independent testing sessions). A cutoff is set at 60s for the intensity level 4, and at 20 s for the intensity level 7). Each intensity stimulation is applied 3 times. The mean time for tail withdrawal is measured.

#### Acetone test

The acetone test is conducted by applying, from underside, a drop of acetone. Mice are placed on a polymethyl methacrylate box with a wire grid floor. The drop is formed with a syringe connected to a thin polyethylene tube to the plantar skin of animal’s hind paw, under the leg in the cutaneous territories of interest. The acetone drop is applied to each animal hind paw twice with an interval of a least one minute, and the duration and the quality of the responses (licking, biting, shacking) are recorded during 1 min.

#### Spared Nerve Injury (SNI) model

The SNI model is applied as initially described by Decosterd and colleagues ^100,121^. The surgery is performed under isoflurane anesthesia (2.5%/100% oxygen mix). The body temperature is maintained during the whole procedure by using a heating pad. Eyes are protected with ophthalmic cream (Viscotear ®). Briefly, the skin and the biceps femoris are incised, exposing the sciatic nerve under a binocular lens. The tibial and common peroneal nerves are ligated with a silk suture 6.0 and transected. The muscle is closed with a 6.0 thread. Then, the skin is closed with a thread-suture (5.0 silk thread). After surgery, the animals are placed in a recovery room (plastic box, at a size comparable to an open field 40 cm x 40 cm, with a thick mattress pad on the bottom and a heating lamp) and a score sheet is filled.

### Patch Clamp Electrophysiology

Adult wildtype or transgenic mice were killed by CO_2_ inhalation, and DRGs from all levels were excised and transferred to DMEM containing 50 μg/ml gentamicin (Sigma-Aldrich, Taufkirchen, Germany). Following treatment with 1 mg/ml collagenase and 0.1 mg/ml protease for 30 min (both from Sigma-Aldrich, Taufkirchen, Germany), ganglia were dissociated using a fire-polished, silicone-coated Pasteur pipette. Isolated cells were transferred onto poly-d-lysine-coated (200 μg/ml, Sigma-Aldrich) coverslips and cultured at 37°C and 5% CO_2_ without NGF. Cells were used for experiments within 24 h after plating. Whole-cell voltage and current-clamp recordings were conducted with a HEKA Electronics USB 10 amplifier combined with Patchmaster software (HEKA Elektronik, Lambrecht, Germany). Currents were filtered at 5 kHz and sampled at 20 kHz. Offline analyses were performed using the Fitmaster software (HEKA Elektronik, Lambrecht, Germany and Origin 2024 (Origin Lap, Northampton, MA, USA). For current clamp recordings, the pipette solution contained 140 mM K-gluconate, 10 mM Na-gluconate, 1 mM MgCl_2_, 10 mM EGTA, 10 mM HEPES, and 3 mM Mg-ATP. pH 7.4 was adjusted with KOH. The external solution contained 140 mM NaCl, 5 mM KCl, 2 mM CaCl_2_, 2 mM MgCl_2_ and 10 mM HEPES. For voltage clamp recordings, the pipette solution contained 140 mM CsCl, 2 mM MgCl2, 5 mM EGTA, 10 mM HEPES and 3 mM Mg-ATP. pH 7.4 was adjusted with CsOH. The extracellular solution contained 140 mM tetraethylammonium chloride, 1 mM MgCl_2_, 5 mM CaCl_2_ and 10 mM HEPES, pH 7.4 was adjusted with tetraethylammonium hydroxide. Patch pipettes were fabricated with borosilicate glass (Science Products) using a conventional puller (PC-100 Puller, Narashige, Japan) and heat-polished to give a pipette resistance of 2-3 MΩ.

### RNA-Seq

To perform RNA-Seq analysis of cultured DRG neurons, we prepared DRGs from 14 male C57BL/6N mice (10 weeks old) per replicate experiment (see Dorsal root ganglion neuron cultures). The isolated cells were seeded in 7 wells of 6-well plates containing 2 ml supplemented NB medium per well and cultured overnight. On the next day, the cell cultures were stimulated with respective compound combinations for 6 h, rinsed with 1 ml fresh medium, and subsequently lysed in 500 µl Nucleozol (Macherey Nagel). RNA isolation was performed according to manufacturer’s instructions (NucleoSpin RNA Set for NucleoZol, Macherey-Nagel). RNA integrity (RIN) was assessed using a TapeStation (Agilent Technologies) and was > 9.1 for all samples.

For transcriptomic analysis of Cacna1c^fl/fl^ and SNS-Cacna1c^-/-^ mice, all DRGs from each mouse (24 weeks old) were collected in 2.0 ml RNAlater (Thermo Fisher Scientific, #AM7020) and incubated overnight at 4°C. RNAlater was then carefully removed, and tissues were stored at -70 °C until further processing. DRGs were lysed in 500 µl NucleoZOL (Macherey-Nagel) using a tissue lyser (Qiagen, 2 min, 50 Hz) with precooled stainless-steel beads (5 mm diameter, #69989, Qiagen). Lysates were centrifuged (30 s, 17,000 g), and total RNA was extracted according to the manufacturer’s protocol (NucleoSpin RNA Set for NucleoZOL, Macherey-Nagel). RIN values were > 8.

Libraries were prepared with the Illumina TruSeq kit according to the standard protocol with Poly A selection. After validation (2200 TapeStation, Agilent Technologies) and quantification (Qubit System; Invitrogen) pools of cDNA libraries were generated. The pools were quantified using the KAPA Library Quantification kit (Peqlab) and the 7900HT Sequence Detection System (Applied Biosystems) and subsequently sequenced on an Illumina HiSeq4000 sequencer using a 2 × 100 bp protocol. Reads were mapped to the murine genome (mm10) and quantified using Salmon ^113^. Data were normalized, and statistics were calculated using DESeq2 ^112^.

### Statistical Analysis

Statistical analyses were performed with Students t-tests, one-, or two-way ANOVA with respective post hoc tests as indicated in the figure legends. *P* < 0.05 was considered statistically significant. Dose-response curves from HCS microscopy were generated using non-linear regression curve-fitting (three parameter, standard Hill slope) with Prism (GraphPad, La Jolla, CA). The parameters of the model (top, bottom, or pEC_50_/pIC_50_ values) were compared using the extra-sum-of-squares F test. High content screening kinetic experiments were analyzed with R using ordinary two-way ANOVA. Bonferroni’s post hoc analysis was applied to determine *P* values of selected pairs defined in a contrast matrix using the R library multcomp. Error bars represent the standard error of the mean (SEM) of 3-5 independent replicate experiments using cells of different animals.

## Supplemental information

Table S1. Excel tables with all detected and differentially expressed genes from the in vitro experiment.

Table S2: Excel tables with all detected and differentially expressed genes from the in vivo experiment.

Table S3: GO analysis of in vivo experiment

