## Supplemental figures for "Ca_V_1.2-dependent excitation-transcription coupling modulates nociception"

### Supplement for manuscript: $Ca_V$1.2-dependent excitation-transcription coupling modulates nociception

### SUPPLEMENTAL INFORMATION

Supplemental figures S1-S10

Table S1. Excel tables with all detected and differentially expressed genes from the in vitro experiment.

Table S2: Excel tables with all detected and differentially expressed genes from the in vivo experiment.

Table S3: GO analysis of in vivo experiment

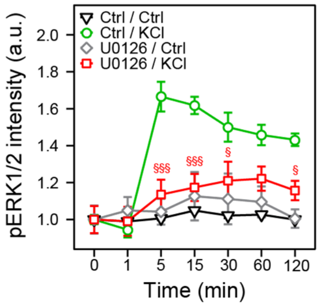


**Figure S1: Depolarization induces phosphorylation of ERK1/2, related to Figure 1. (A)** Effect of the MEK inhibitor U0126 on KCl-induced (40 mM) ERK1/2 phosphorylation. Data represent means ± SEM; n = 3-4 experiments; >2000 neurons/condition; two-way ANOVA with Bonferroni's test. §P<0.05; §§P<0.01; §§§P<0.001 indicate significance levels between KCl-induced pERK1/2 signals in the absence or presence of the inhibitor.

**A**

| 21822 genes q < 0.05 | KCl regulated | | | BayK regulated | | | VP regulated | | |
|---|---|---|---|---|---|---|---|---|---|
| | KCl10 vs. Ctrl A_B | KCl40 vs. Ctrl A_C | KCl40 vs. KCl10 B_C | BayK vs. Ctrl A_D | KCl10+BayK vs. Ctrl A_E | KCl10+BayK vs. KCl10 B_E | Ctrl vs. VP A_F | KCl40+VP vs. Ctrl A_G | KCl40 vs. KCl40+VP C_G |
| **FC > 0** | 833 | 3391 | 2464 | 56 | 1066 | 34 | 329 | 3503 | 473 |
| **FC < 0** | 831 | 4000 | 3059 | 74 | 1392 | 23 | 159 | 3968 | 184 |
| **FC > 1.5** | 51 | 667 | 461 | 16 | 116 | 11 | 20 | 777 | 23 |
| **FC < 0.67** | 77 | 896 | 542 | 6 | 122 | 3 | 11 | 980 | 30 |
| **FC > 2** | 6 | 201 | 115 | 5 | 17 | 3 | 1 | 228 | 2 |
| **FC < 0.5** | 4 | 171 | 77 | 0 | 11 | 1 | 0 | 223 | 6 |

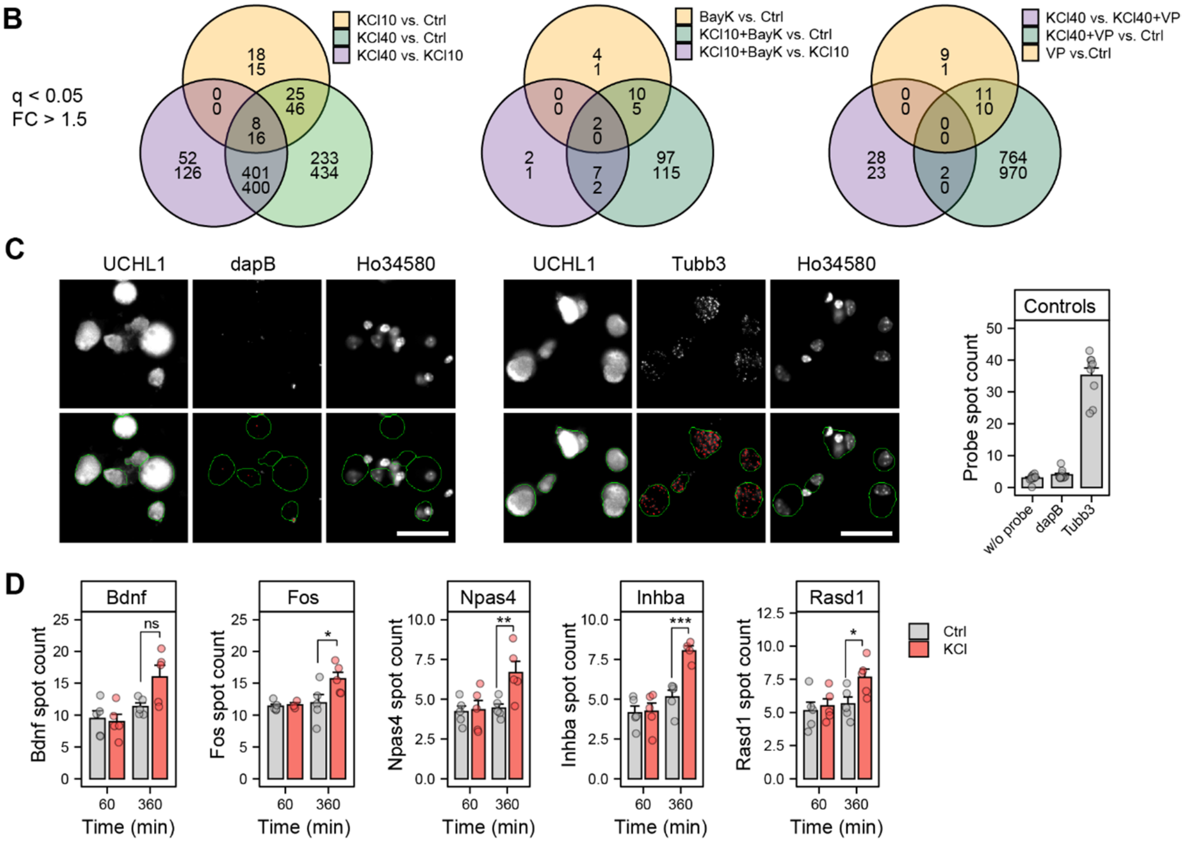


**Figure S2: Depolarization leads to Cav1-dependent regulation of gene expression in sensory neurons, related to Figure 2. (A)** Numbers of differentially expressed genes identified by RNA-Seq at different fold-change cut-offs. **(B)** Overlap of differentially expressed genes KCl, BayK8644 (BayK), and verapamil (VP). **(C)** Controls showing the detection of βIII-tubulin (positive control) versus the bacterial gene dapB (negative control) by single molecule Fluorescence In Situ Hybridization (smFISH). The sensory neurons were detected using a UCHL1-specific antibody, nuclei were labeled with Hoechst 34580. **(D)** Quantification of *Bdnf*, *Fos*, *Npas4*, *Inhba*, and *Rasd1* mRNA expression by smFISH in UCHL1-labeled sensory neurons after KCl-depolarization (40 mM) for 60 or 360 mins. Data represent means ± SEM; n = 4-5 experiments; >3000 neurons/condition; two-way ANOVA with Bonferroni's test. *$p$<0.05; **$p$<0.01; ***$p$<0.001.

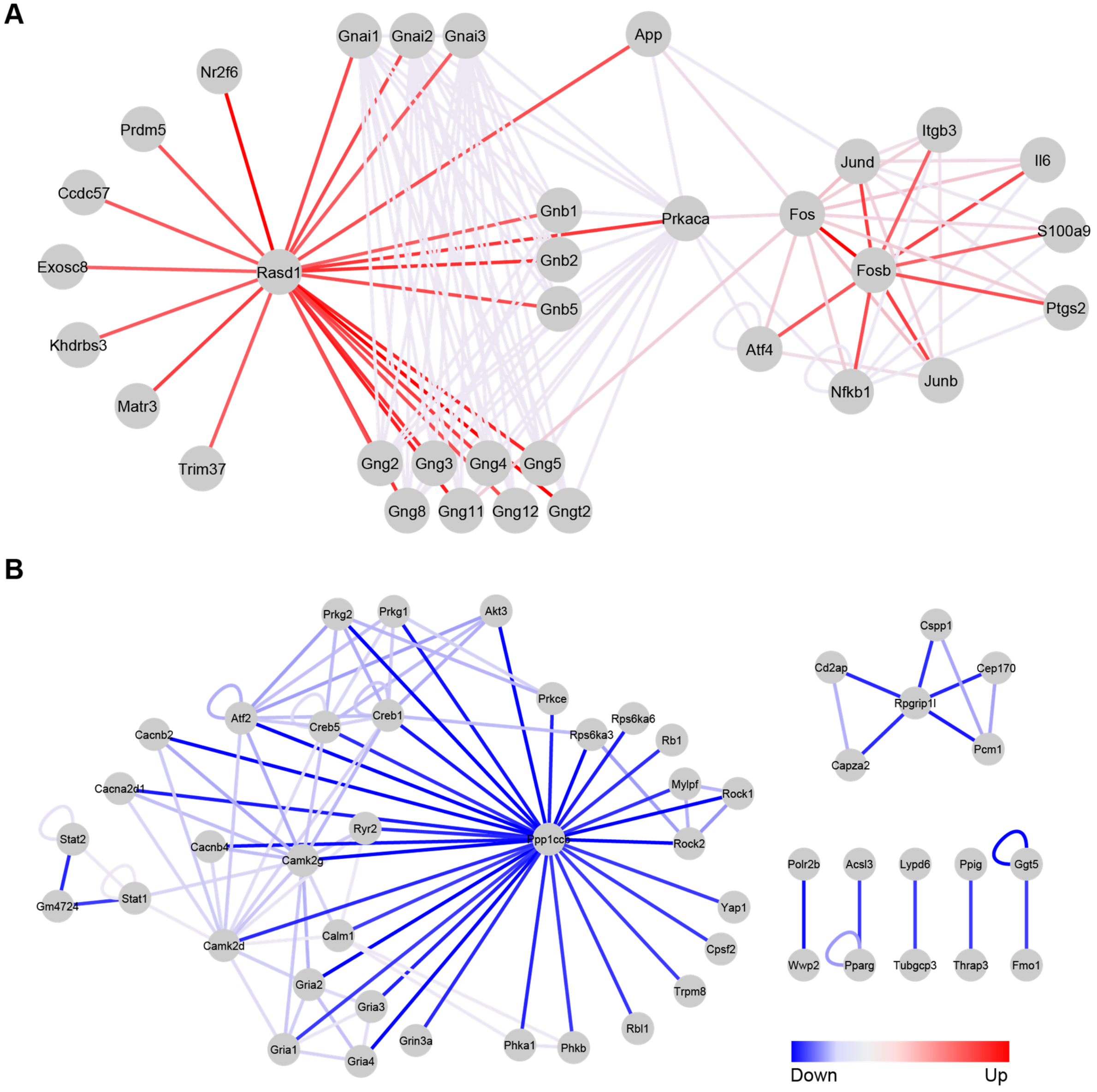


**Figure S3: Protein modules (neighborhoods) identified by calculating conditional interactions scores for all protein pairs in the mouse network and then identifying those interactions that are most strongly altered in a coherent manner following BayK treatment compared to KCl(10) alone. These subnetworks were obtained by selecting the proteins participating in the most altered interactions and extracting the complete subnetwork, including the selected interactions, and other interactions among selected proteins, related to Figure 2. (A)** The most increased interactions involve Rasd1 and Fosb and form an interconnected network. **(B)** The decreased interactions are centered on the regulatory and catalytic subunit of protein phosphatase 1.

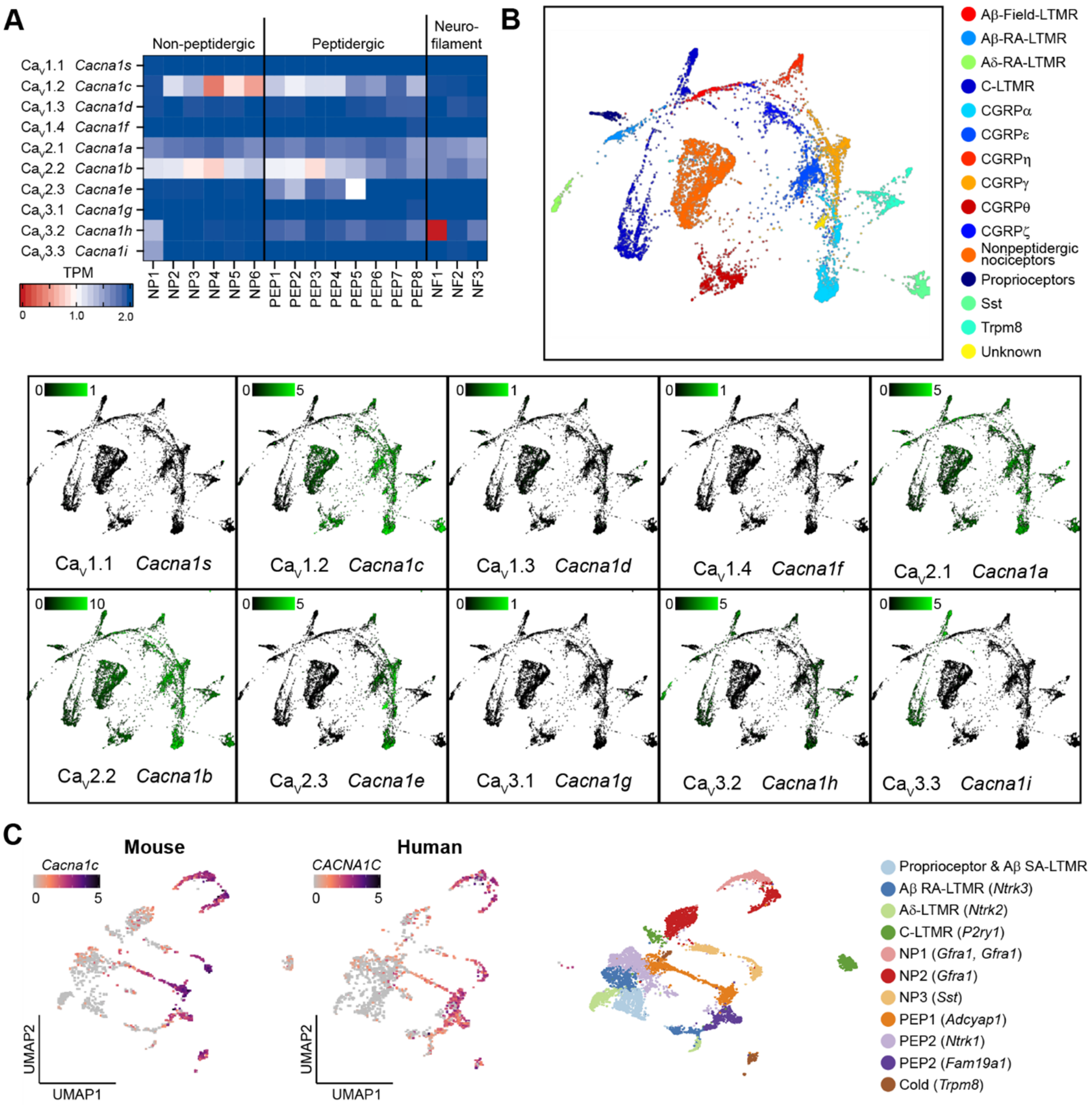


**Figure S4: Cav1.2 is predominantly expressed nociceptive DRG neurons, related to Figure 3.** **(A-B)** Expression pattern of VGCC α subunits in adult mouse DRG neuron subgroups determined by single-cell RNA-seq by Zeisel et al. {Zeisel, 2018 #1920} (A) or Sharma et al. {Sharma, 2020 #2090} **(B)** **(C)** Cross-species comparison of single cell RNA-Seq data from mouse (left) or human (middle) DRG neurons supporting that the Cacna1c expression pattern is conserved {Jung, 2023 #2371}.

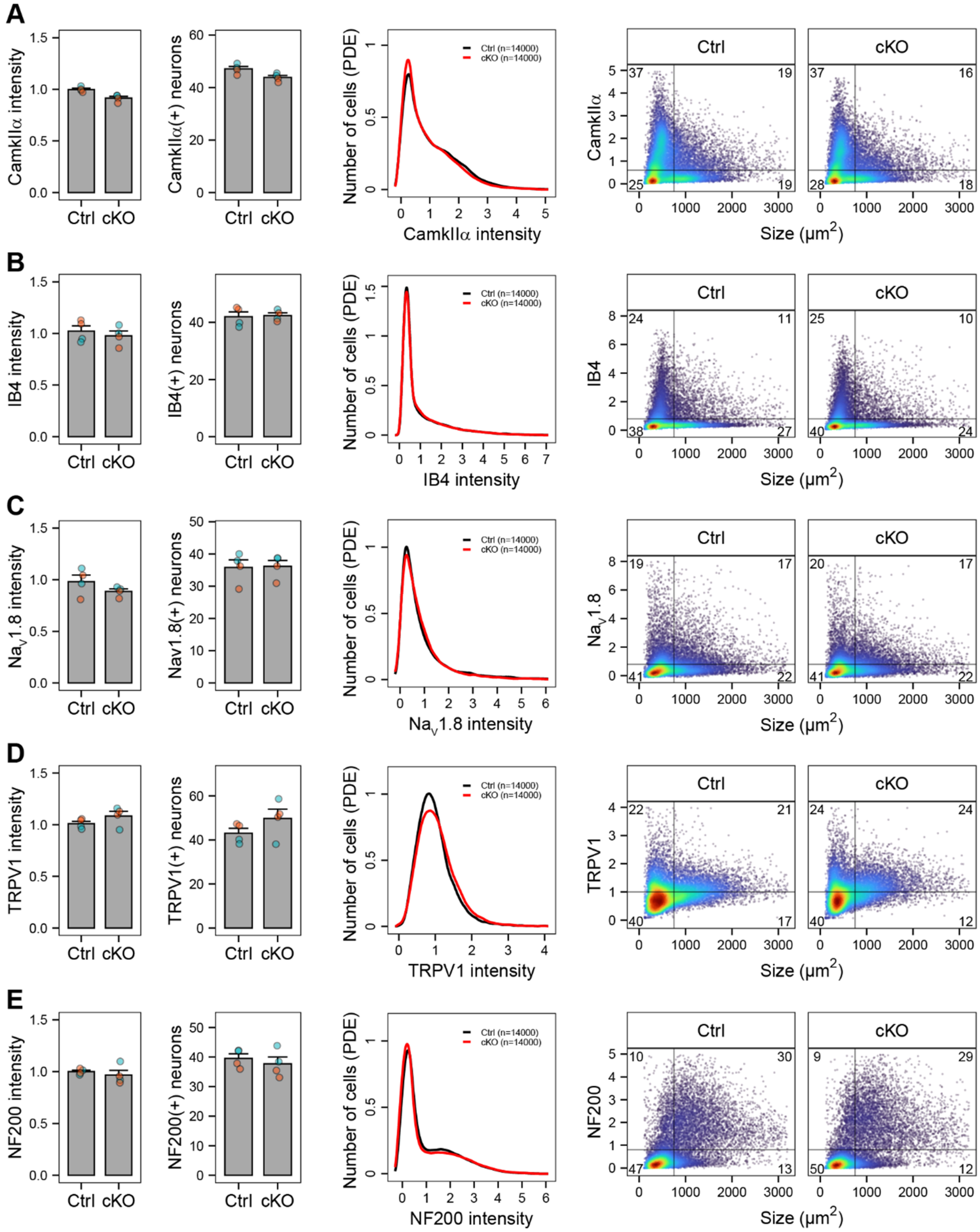

**Figure S5: Cav1.2-deficiency does not affect the composition of neuronal DRG subgroups, related to Figure 3.** Expression pattern of marker proteins for subgroups of sensory neurons expressing CaMKIIα **(A)**, IB4 **(B)**, Nav1.8 **(C)**, TRPV1 **(D)**, and NF200 **(E)**.

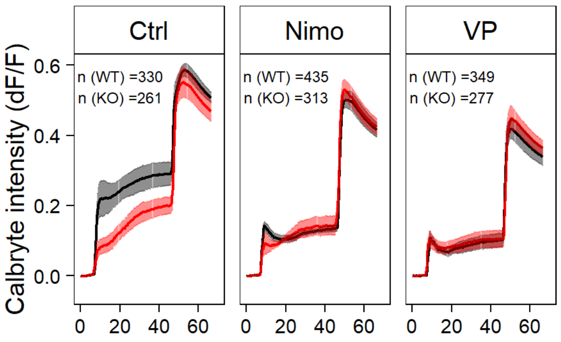


**Figure S6: Cav1.2-deficiency decreases calcium influx after depolarization of DRG neurons, related to Figure 4 (A)** Mean traces after depolarization with a low dose of KCl (10 mM) in the absence or presence of the Cav antagonists nimodipine (Nimo, 10 µM) or verapamil (VP, 100 µM). The plots for Ctrl and Nimo are also shown in Figure 4.

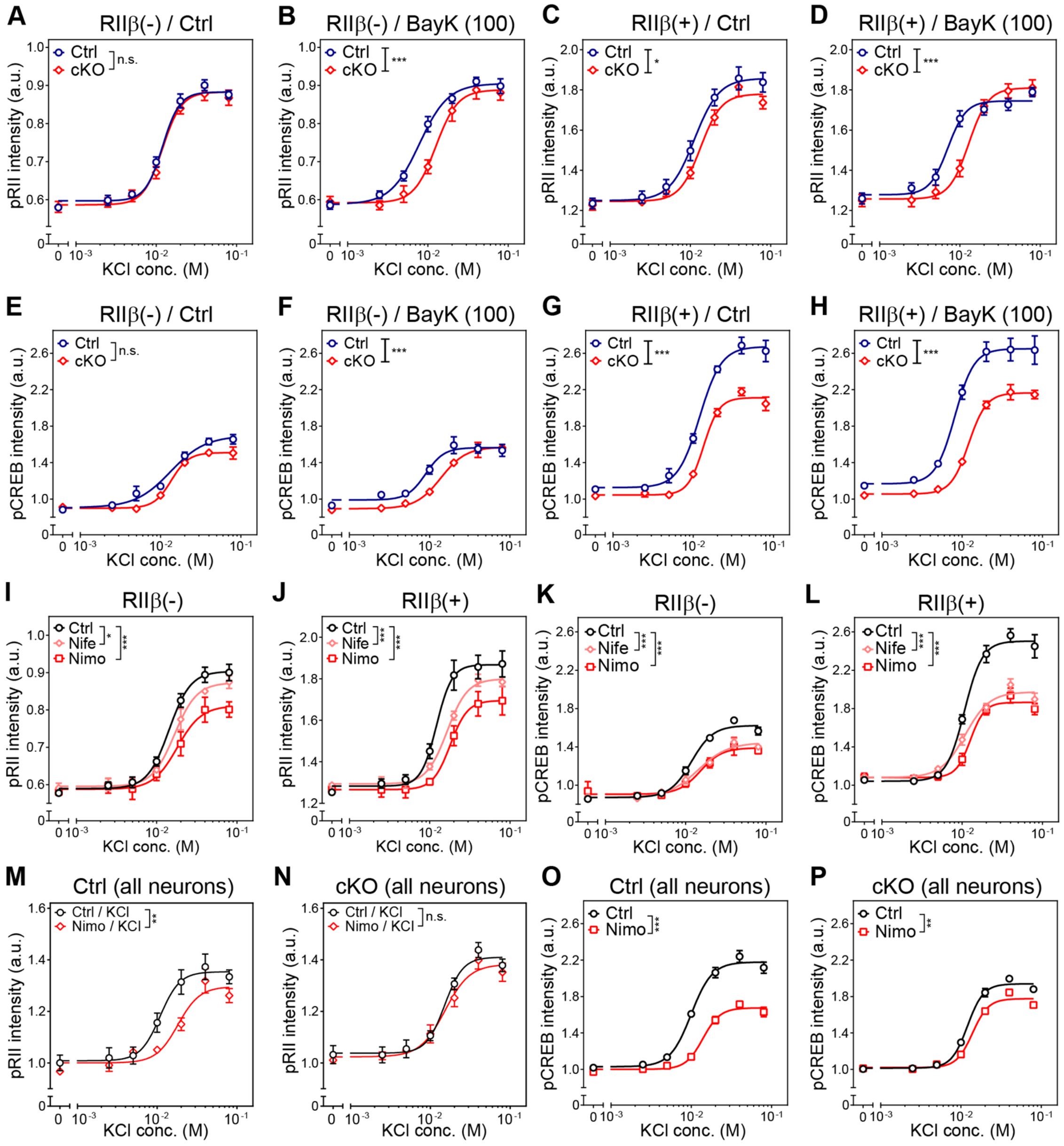


**Figure S7. Cav1.2-deficiency decreases depolarization-induced PKA-II and CREB activity, related to Figure 5. (A-D)** Dose-response curves of pRII intensities in RIIβ(-) and RIIβ(+) DRG neurons of Cacna1c$^{fl/fl}$ and SNS-Cacna1c$^{-/-}$ mice depolarized with KCl (0-80 mM) for 3 min. Neurons in B,D were preincubated with BayK (10, 100 nM) for 10 min. **(E-H)** Dose-response curves of pCREB intensities in RIIβ(-) vs. RIIβ(+) DRG neurons. **(I-L)** Dose-response curves showing the effect nimodipine or nifedipine (10 µM, 10 min) on pRII (I-J) or pCREB (K-L) signals induced by KCl-depolarization. **(M-P)** Dose-response curves of pRII (M-N) or pCREB (O-P) intensities in DRG neurons of Cacna1c$^{fl/fl}$ or SNS-Cacna1c$^{-/-}$ mice in the absence or presence of nimodipine (10 µM, 10 min). Data are means ± SEM; *n* = 3-4 experiments with n > 2000 neurons/condition from female mice; extra-sum-of-squares F test; *$p < 0.05$; **$p < 0.01$; ***$p < 0.001$ indicate significance levels between genotypes or stimulated conditions.

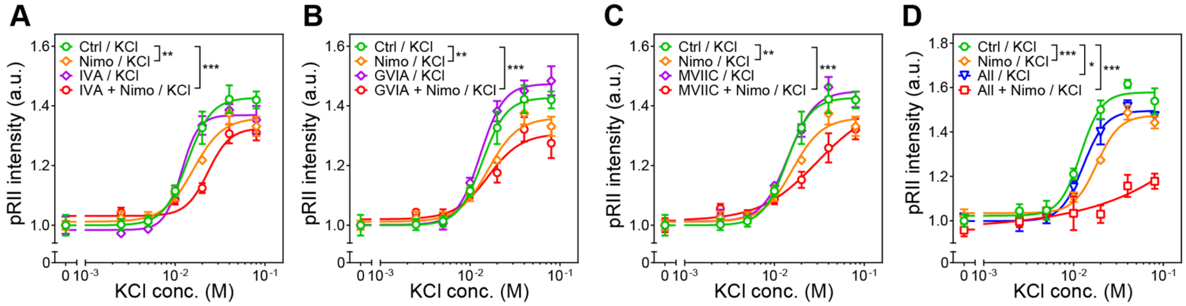


**Figure S8. Cav1.2-deficiency decreases depolarization-induced PKA-II and CREB activity, related to Figure 5. (A-D)** Dose-response curves of pRII intensities in DRG neurons depolarized with KCl (0-80 mM) for 3 min. Neurons in were preincubated for 10 min with nimodipine and/or the Cav2 inhibitors ω-agatoxin IVA (1 µM, Cav2.1, A), ω-conotoxin GVIA (3 µM, Cav2.2, B), and ω-conotoxin MVIIC (1 µM, Cav2, C). In (D), all Cav2 inhibitors (All) were combined and tested in the absence or presence of nimodipine.

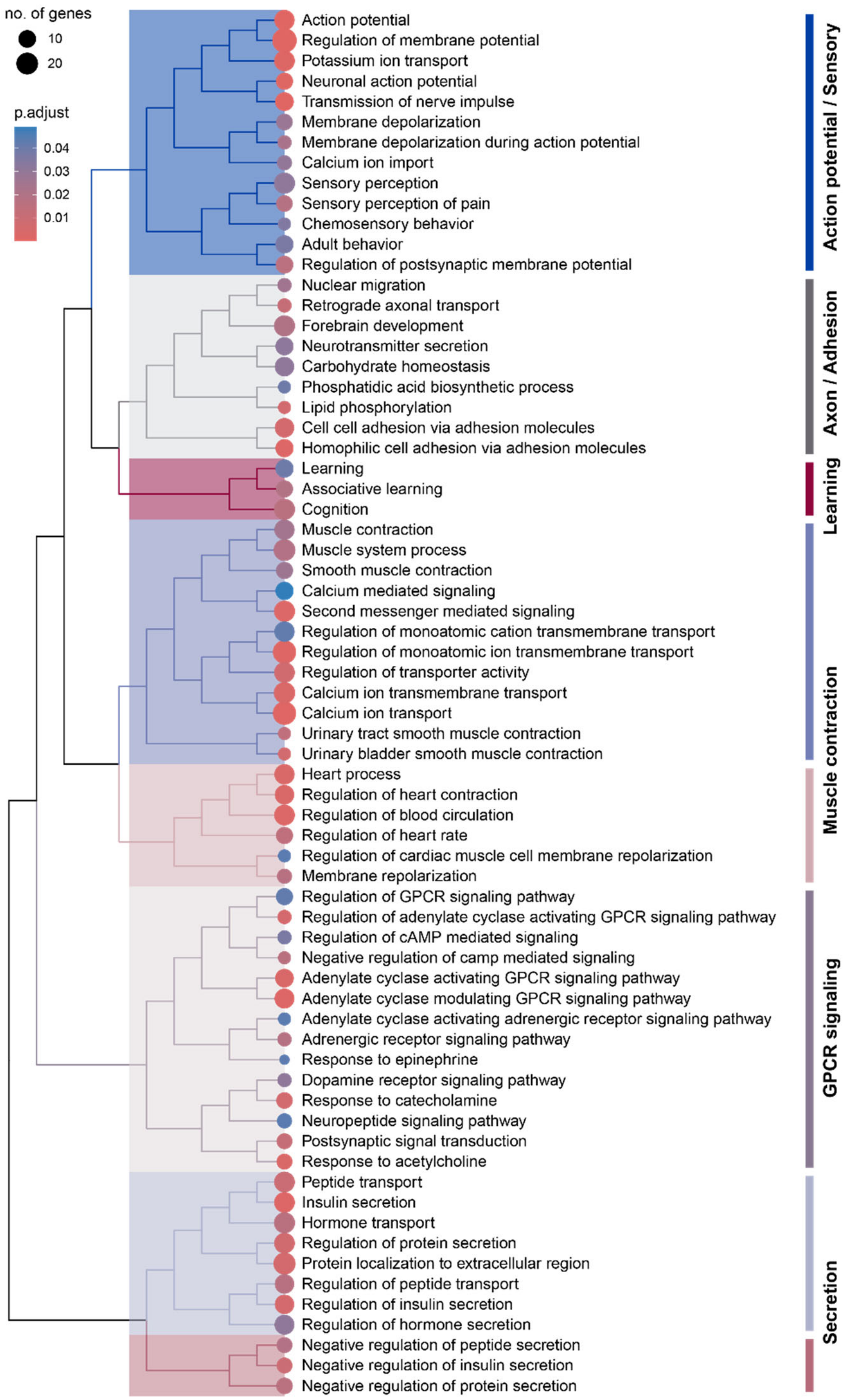


**Figure S9. Cav1.2-dependent E-T coupling induces transcriptional regulation *in vivo*, related to Figure 6.** Hierarchical clustering of the enriched terms among differentially expressed genes in DRGs of Cacna1c$^{fl/fl}$ versus SNS-Cacna1c$^{-/-}$ mice. The approach relies on the pairwise similarities of the enriched terms calculated by the pairwise_termsim() function, which by default using Jaccard's similarity index (JC).

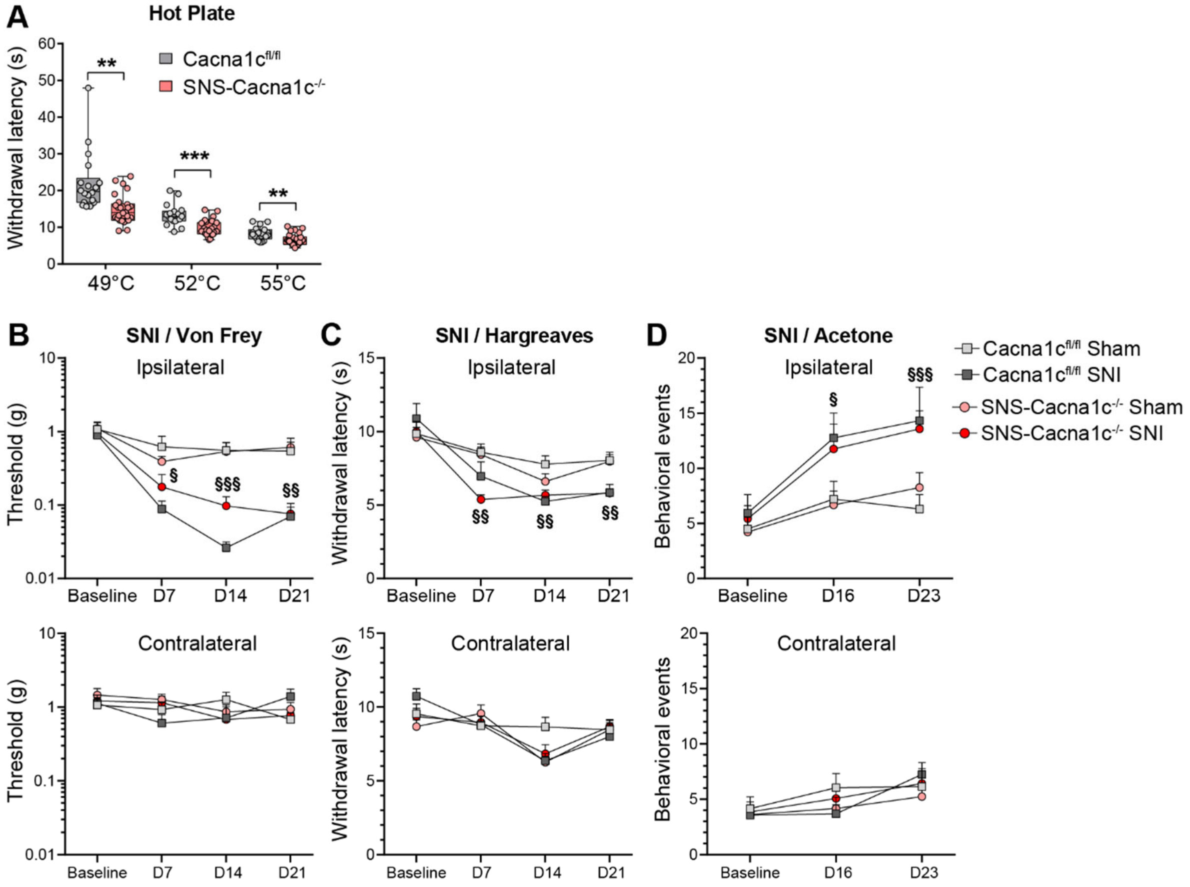


**Figure S10. SNS-Cacna1c$^{-/-}$ mice show increased heat nociception and CFA-induced inflammatory pain, related to Figure 8. (A)** Acute pain responses to noxious heat determined with the hot plate test (n = 18 Cacna1c$^{fl/fl}$ and 26 SNS-Cacna1c$^{-/-}$ male and female mice, 2-way ANOVA with Tukey's post hoc test). **(B)** Mechanical sensitivity measured by Von Frey filaments at the ipsilateral (top) or contralateral (bottom) hindpaw after SNI surgery (n = 10 Cacna1c$^{fl/fl}$ Sham, n = 8 Cacna1c$^{fl/fl}$ SNI, n = 12 SNS-Cacna1c$^{-/-}$ Sham, n = 14 SNS-Cacna1c$^{-/-}$ SNI, male and female mice, 2-way ANOVA with Tukey's post-hoc test. § $p < 0.05$, §§ $p < 0.01$, §§§ $p < 0.001$ between Sham and SNI). **(C)** Thermal heat sensitivity measured by Hargreaves method after SNI surgery. **(D)** Thermal cold sensitivity measured using the Acetone test after SNI surgery.
